# KRAS interaction with RAF1 RAS-binding domain and cysteine-rich domain provides insights into RAS-mediated RAF activation

**DOI:** 10.1101/2020.07.31.231134

**Authors:** Timothy H. Tran, Albert H. Chan, Lucy C. Young, Lakshman Bindu, Chris Neale, Simon Messing, Srisathiyanarayanan Dharmaiah, Troy Taylor, John-Paul Denson, Dominic Esposito, Dwight V. Nissley, Andrew G. Stephen, Frank McCormick, Dhirendra K. Simanshu

**Author notes:** These authors contributed equally to this work (Timothy H. Tran, Albert H. Chan, Lucy C. Young). To whom correspondence should be addressed: Dhirendra Simanshu), Frank McCormick).

## Abstract

A vital first step of RAF activation involves binding to active RAS, resulting in the recruitment of RAF to the plasma membrane. To understand the molecular details of RAS-RAF interaction, we solved crystal structures of wild-type and oncogenic mutants of KRAS complexed with the RAS-binding domain (RBD) and the membrane-interacting cysteine-rich domain (CRD) from the N-terminal regulatory region of RAF1. Our structures revealed that RBD and CRD interact with each other to form one structural entity in which both RBD and CRD interact extensively with KRAS. Mutation at the KRAS-CRD interface resulted in a significant reduction in RAF1 activation despite only a modest decrease in binding affinity. Combining our structures and published data, we provide a model of RAS-RAF complexation at the membrane, and molecular insights into RAS-RAF interaction during the process of RAS-mediated RAF activation.

## INTRODUCTION

The mitogen-activated protein kinases (MAPK) pathway, also known as the RAS-RAF-MEK-ERK signaling pathway, regulates diverse cellular processes including cell proliferation, differentiation, and survival via phosphorylation cascades in all eukaryotic cells. The interaction of membranebound active KRAS (GTP-bound) with RAF is a key step in the activation of this pathway^1,2^. Activated RAS recruits RAF kinases to the membrane where RAF dimerizes and becomes active. The activated RAF kinases then phosphorylate and activate MEK kinases, which in turn phosphorylate and activate ERK kinases. Finally, activated ERK kinases are translocated into the nucleus, where they phosphorylate multiple substrates that regulate vital cellular processes. RAS mutations resulting in constitutive activation of the MAPK pathway are the most common mutations observed in human cancers. Taking advantage of this, therapeutic strategies that aim to disrupt RAS-RAF interaction are being developed as drug candidates against RAS-driven cancers^3^.

The mammalian RAF family of enzymes comprises three evolutionarily conserved cytosolic serine/threonine kinases (ARAF, BRAF, and RAF1, also known as CRAF), which share three conserved regions, CR1, CR2, and CR3^4^ (**Fig. 1a**). The N-terminal CR1 region contains the RAS-binding domain (RBD) and the cysteine-rich domain (CRD), the CR2 region contains a 14-3-3 recognition site, whereas the C-terminal CR3 region includes the kinase domain and a second binding site for 14-3-3 proteins. The 14-3-3 proteins are a family of highly conserved dimeric proteins that bind to specific phosphorylated serines or threonines present in the conserved sequence motifs. Prior to RAF activation, a 14-3-3 dimer binds to phosphorylated serines present in CR2 and CR3, keeping RAF in an autoinhibited state. A recent cryoEM structure of this state revealed that the CRD interacts with both 14-3-3 and the kinase domain of RAF in a manner that precludes CRD interaction with the membrane^5^.

**Figure 1:**
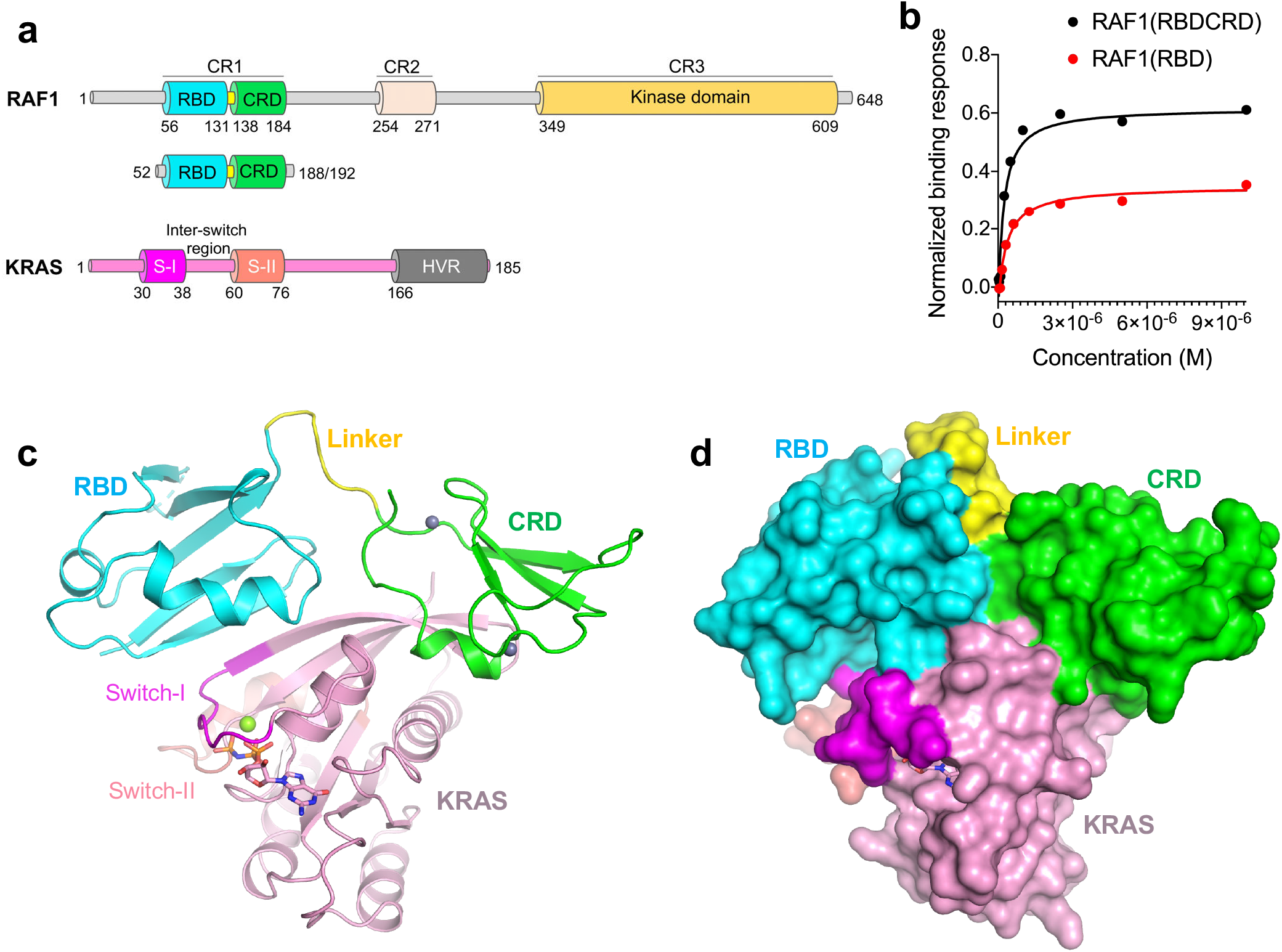
Structure of the KRAS-RAF1(RBDCRD) complex and SPR analysis. **(a)** Domain architecture of RAF1 and KRAS. Key domains and their boundaries are shown. The RAF1 construct containing RBD and CRD regions used in this study is shown below the full-length RAF1. The G-domain of KRAS4b(1-169) used in this study contains switch-I (S-I) and switch-II (S-II) regions with the inter-switch region present between them. **(b)** Steady-state binding isotherms derived from the SPR data used for measuring the binding affinities of RAF1(RBDCRD) and RAF1(RBD) to KRAS-GMPPNP. **(c, d)** The overall structure of the complex formed by GMPPNP-bound KRAS and RAF1(RBDCRD) in (c) cartoon, and (d) surface representations. GMPPNP is shown as sticks, and Mg^2+^ (green) and Zn^2+^ (gray) are shown as spheres. KRAS is colored pink with switch-I and II regions highlighted in magenta and salmon, respectively. RBD, CRD, and the linker present between them are colored cyan, green and yellow, respectively.

Autoinhibited RAF can be activated by membrane-anchored active RAS, which binds to the RBD with high affinity (nanomolar). Following recruitment of RAF to the membrane, the CRD forms contacts with phospholipids, disrupting the autoinhibited state. Based on mutagenesis and NMR experiments, CRD residues involved in membrane interaction have been proposed to be located in the two hydrophobic loops surrounded by basic residues^6–8^. In addition to the known RBD-RAS interactions, CRD has been reported as a second RAS-binding domain of RAF^9–13^ that can bind to RAS independently of RBD with weaker (micromolar) affinity^14^. Interaction between CRD and RAS is critical for activation of RAF^12,15–18^. RAS residues N26 and V45, and CRD residues S177, T182 and M183 play an essential role in RAS-CRD interaction^9,12^. NMR and mutagenesis studies also suggested that CRD may interact with RAS residues from 23 to 30^11^, residues in the switch-II region (G60 and Y64)^10^, as well as the farnesyl group of RAS^11,14^. However, the importance of the farnesyl-CRD interaction at the membrane remains unclear. The CRD appears to play critical roles in maintaining RAF kinase in the inactivate state, and in the process of RAS-dependent RAF kinase activation through direct interactions with the plasma membrane and RAS itself.

Although crystal structures of HRAS and Rap1 in complex with RBD, and an NMR structure of an isolated CRD have been solved^19–21^, no experimental structure of the RAS-RBDCRD or RAS-CRD complex is currently available. Recent reports of the cryoEM structures of full-length BRAF in complex with 14-3-3 and MEK have provided new insights into the autoinhibited and active RAF complexes. However, none of these structures showed the first half of BRAF in its entirety^5,22^, presumably because conserved regions CR1 and CR2 are too flexible in the absence of RAS. While progress has been made in understanding the role of RAS in RAF activation, the precise molecular details underlying the interaction of RAS and RAF via the formation of a RAS-RBDCRD complex at the cell membrane, and the role of CRD in stabilizing the active RAS-RAF complex remain unclear. Here, we present crystal structures of wild-type and oncogenic mutants of KRAS in complex with the RBD and CRD present at the N-terminal regulatory region of RAF1. The structures provide detailed insights into the KRAS-CRD interaction interface and show how RBD and CRD are arranged with respect to one another. They also extend the footprint of RAF binding to RAS beyond the classical, highly conserved effector/RBD binding region in the switch I, to residues 23, 24 and 26, the inter-switch region (42-45) and even residues 149, 153 and 157 in helix a5. The inter-switch region differs significantly amongst RAS-related proteins in the RRAS, RIT and RAP families and may explain why all of these proteins can bind RAF kinases, but only HRAS, KRAS and NRAS proteins activate RAF kinases efficiently.

Structure-based mutagenesis studies identified the role of various interfacial residues in KRAS-RBDCRD interaction as well as RAS-mediated RAF activation. Based on published mutagenesis, NMR and molecular dynamic (MD) simulation studies, we suggest how the KRAS-RAF1(RBDCRD) complex (hereinafter referred to as KRAS-RBDCRD) might interact with the membrane. Combining the new structural insights obtained from our KRAS-RBDCRD structures and the recently solved autoinhibited state of BRAF in complex with 14-3-3 and MEK, we show molecular details of RAS-RAF interaction during the process of RAS-mediated RAF activation.

## RESULTS

### Structures of KRAS in complex with two RAS-binding domains of RAF1

To understand how the tandem domains, RBD and CRD, in the N-terminal regulatory part of RAF1 interact with KRAS, and their roles in RAS-mediated RAF activation, we attempted to crystallize and solve the structure of KRAS in complex with the RBDCRD region of RAF1 (**Fig. 1a**). We made multiple constructs of RAF1(RBDCRD). Among these, two constructs with residues ranging from 52-188 and 52-192 yielded soluble and stable proteins after expression and purification. In addition, we expressed and purified RAF1(RBD) (residues 52-131). Measurement of binding affinities by surface plasmon resonance (SPR) showed high-affinity interaction (K_D_ ~371 nM) between KRAS and RAF1(RBD), as shown previously^23^. The presence of CRD in the RAF1(RBDCRD) resulted in approximately a two-fold increase in the binding affinity with KRAS (K_D_ ~152 nM), suggesting that CRD also plays a role in KRAS-RBDCRD interaction (**Fig. 1b, Supplementary Fig. 1a**).

We crystallized the GTPase domain of KRAS (residues 1-169) in complex with RAF1(RBDCRD) (residues 52-188) and solved the structures in two different crystal forms, I and II, at a resolution of 2.15 Å and 2.50 Å, respectively (**Table 1, Supplementary Table 1**). Despite having different crystal packing interactions, the structures of both crystal forms are very similar, with r.m.s. deviation of 0.393 Å for Cα atoms or 0.445 Å for all atoms (**Supplementary Fig. 1b**). Since crystal form I was solved at a higher resolution, subsequent analyses are based on form I unless stated otherwise. The overall structure shows that the two tandem domains of the RAF1 CR1 region, RBD and CRD, directly contact one another to form one structural entity, and RBD as well as CRD interact with KRAS to a similar extent (**Fig. 1c, d**). The structure of the KRAS-RBDCRD complex allows us to gain insights into the different interaction interfaces such as KRAS-RBD, RBD-CRD, and KRAS-CRD. Similar to what have been observed in the HRAS-RAF1(RBD) and Rap1-RAF1(RBD) structures^19,21^, the KRAS-RBD interaction interface is mainly formed by KRAS residues present in the switch-I region, where RBD and KRAS interact mainly via β-strands and form an extended β-sheet structure. Interestingly, the CRD interaction with KRAS does not involve the switch regions. Instead, the KRAS-CRD interface is formed by KRAS residues present in the inter-switch region and the C-terminal helix α5 (**Supplementary Fig. 1c**). Like the KRAS-RBD interface, KRAS residues involved at the KRAS-CRD interaction interface are conserved across all four RAS isoforms (**Supplementary Fig. 1c**). It has been suggested that RBD and CRD form two independent globular domains, where RBD’s role is to interact with KRAS, and CRD helps anchor RAF1 to the membrane. Our results suggest that CRD plays not only an important role in anchoring RAF1 to the membrane, but also a direct role in enhancing RAS-RAF interaction.

**Table 1.**
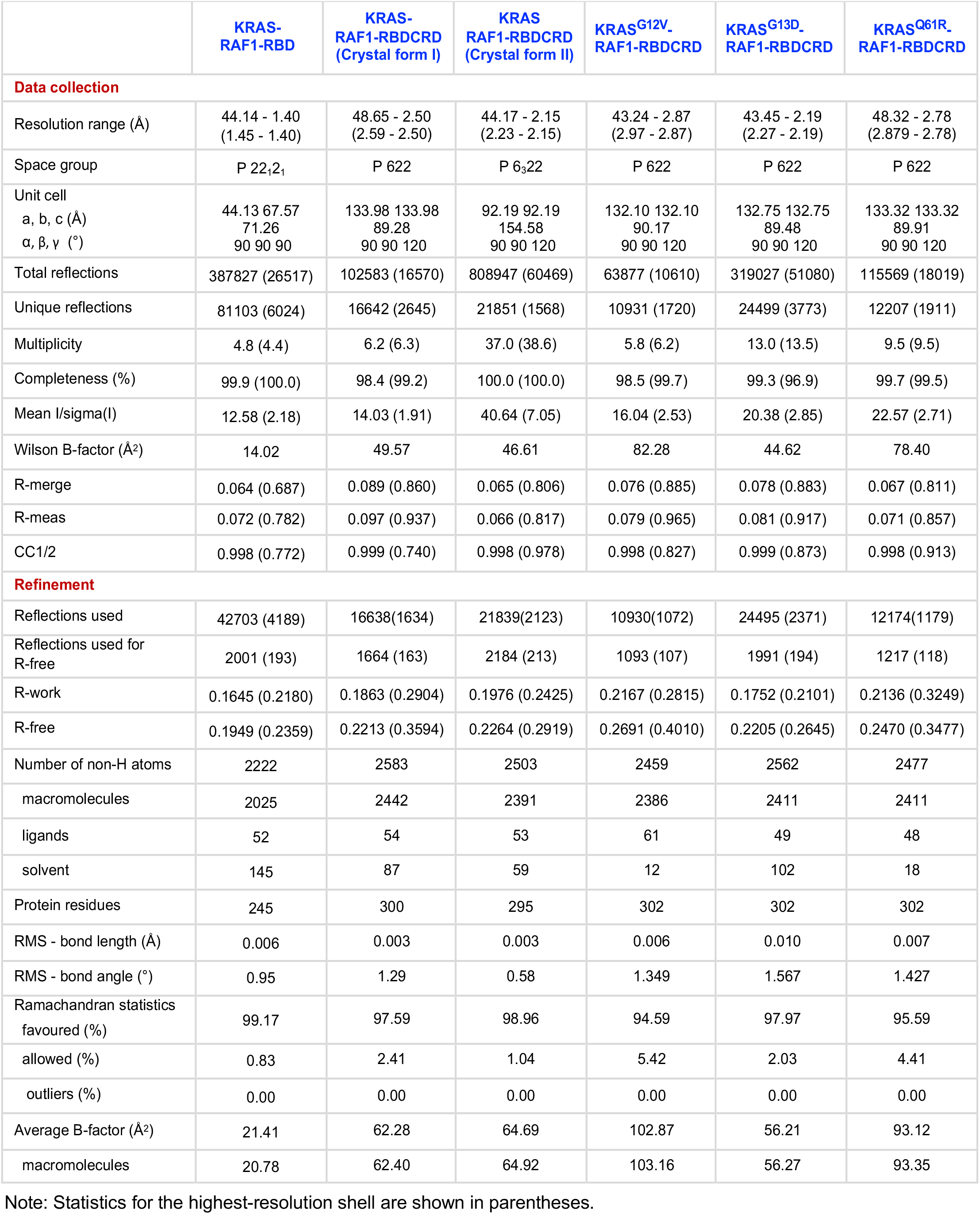
Crystallographic data collection and refinement statistics.

### KRAS interaction with RBD plays a major role in the high-affinity KRAS-RAF1 complex formation

To understand what impact the presence of CRD may have on the KRAS-RBD interaction, we solved the structure of KRAS in complex with RAF1(RBD) and compared it with the KRAS-RBDCRD structure (**Table 1, Supplementary Table 1**). Structural superposition of KRAS-RAF1(RBD) (hereinafter referred to as KRAS-RBD) with KRAS-RBDCRD using KRAS residues shows no significant differences at the KRAS-RBD interaction interface in the two structures (**Fig. 2a**). However, RBD residues that are located away from the KRAS-RBD interface show significant conformational changes. Structural analysis shows that the RBD domain in the KRAS-RBDCRD structure undergoes a rotation of 9.3° and a displacement of 0.9 Å compared to that of the KRAS-RBD structure (**Fig. 2b**). This observation suggests that KRAS-RBD complex is likely to be flexible in the absence of CRD, and this rotation of RBD may be needed to properly orient CRD to stabilize the KRAS-RBDCRD interaction. This is consistent with an MD simulation study which suggested that CRD reduces the fluctuations of the KRAS-RBD complex at the membrane and enhances its binding affinity^24^.

**Figure 2:**
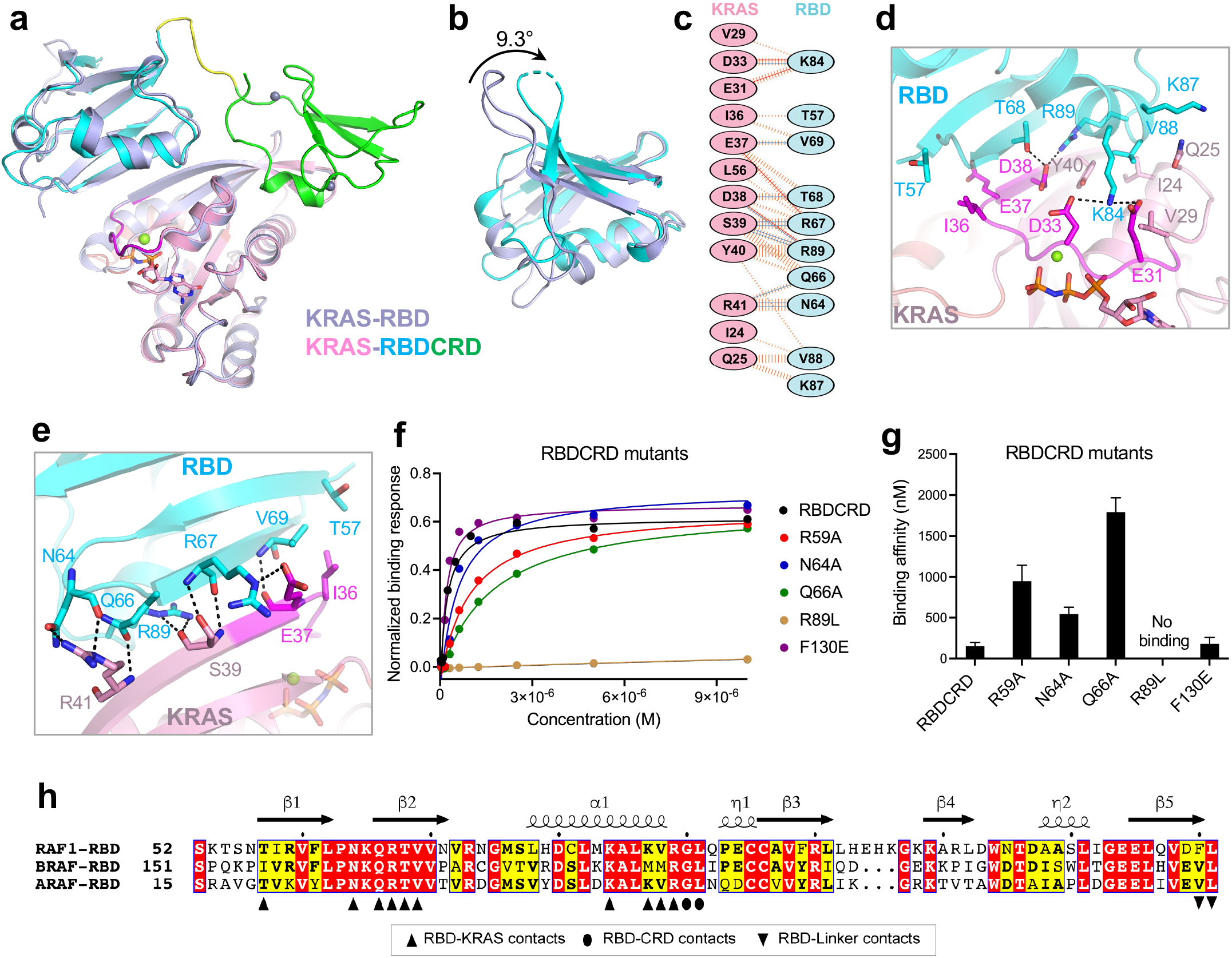
Structural and mutational analyses of the KRAS-RBD interactions in the KRAS-RBDCRD complex. **(a)** Superposition of KRAS-RBD structure (light blue) with KRAS-RBDCRD structure shows conformational differences in RBD. The structural superposition was carried out by aligning atoms in KRAS in both structures. **(b)** Enlarged view of RBD showing conformational differences that occur when it binds to KRAS as RBD vs. RBDCRD. **(c)** Schematic representation of the KRAS-RBD interaction interface, as identified by PDBSum (http://www.ebi.ac.uk/pdbsum/). The interactions are colored using the following notations: hydrogen bonds – solid blue lines, salt bridges – solid red lines, non-bonded contacts – striped orange lines (width of the striped line is proportional to the number of atomic contacts). **(d, e)** Enlarged view of the KRAS-RBD interaction interface formed by residues present in the (d) switch-I region (magenta), and (e) β2 strand (residues present at the end of the switch-I region) of KRAS. The nucleotide GMPPNP and residues that participate in the protein-protein interaction are shown in stick representation. Dashed black lines indicate intermolecular hydrogen bonds and salt bridges. **(f)** Steady-state binding isotherms derived from the SPR data for different point mutants of RBDCRD binding to KRAS-GMPPNP. **(g)** A bar graph showing binding affinity (K_D_) obtained using the SPR data shown in (f) for point mutants of RBDCRD located at the KRAS-RBD interface. **(h)** Amino acid sequence alignment of residues present in the RAS-binding domain (RBD) of human RAF1, BRAF, and ARAF. Fully and partially conserved residues among the RAF isoforms are highlighted in red and yellow, respectively. The secondary structure of RAF1(RBD) is shown above the alignment. The RBD residues that are involved in the interaction with KRAS, CRD, and the linker region are indicated below the alignment with upright triangles, ovals and inverted triangles, respectively.

The KRAS-RBD interface is mainly composed of hydrogen bonds and electrostatic interactions, where acidic residues located in and around the switch-I region (residues from I24 to R41) of KRAS interact with basic residues present in the β2 strand (N64-V69) and a1 helix (K84-R89) of RBD (**Fig. 2c-e, Supplementary Fig. 2a-c**). A total of eight hydrogen bonds and four salt-bridges at the KRAS-RBD interface contribute to the nanomolar affinity between KRAS and RBD (**Fig. 2c)**. Most of these interactions are observed in both KRAS-RBD and KRAS-RBDRCD structures (**Supplementary Table 2 and 3**). In addition, crystal form II of the KRAS-RBDRCD structure contains an extra salt-bridge between RBD R59 and KRAS E37. These interactions resemble the protein-protein interactions observed in the previously solved structures of HRAS-RAF1(RBD) and Rap1-RAF1(RBD)^19,21^.

To understand the role of RBD residues in the KRAS-RBDCRD interaction interface, we mutated five RBDCRD residues (R59A, N64A, Q66A, R89L, and F130E) and examined their binding affinity to active WT-KRAS (GMPPNP-bound) using an SPR assay (**Fig. 2f, g, Supplementary Fig. 2d**). Analysis of the binding affinity shows that R59A, N64A and Q66A mutants in RBDCRD resulted in a 3-12-fold reduction in the binding affinity with KRAS, whereas mutation of R89L resulted in almost complete loss of interaction between RBDCRD and KRAS. These results are consistent with the previous observation that polar and charged residues present on RBD play a key role in forming high-affinity interactions between KRAS and RBD^25^. Conversely, point mutation of a hydrophobic amino acid F130 present at the RBD-linker interface showed no impact on KRAS-RBDCRD interaction.

Amino acid sequence alignment shows that seven out of ten RBD residues involved at the KRAS-RBD interaction interface are conserved across all three RAF isoforms (**Fig. 2h**). The other three residues also exhibit identical amino acids at these positions in two of the three RAF isoforms, suggesting that all three RAF isoforms interact with RAS proteins in a similar manner. RAF1(RBD) residues G90 and L91 form interdomain interactions with CRD and are also conserved across all three RAF isoforms.

### A short linker present between RBD and CRD brings the two domains together

In the KRAS-RBDCRD structure, RBD and CRD are connected by a short (residue 132-137) linker, which brings these two domains together with interdomain contacts, resulting in an extended structure (**Fig. 3a**, upper panel). Mapping the KRAS interaction interface on the RBDCRD surface shows that both RBD and CRD contribute significantly to RAF1 interaction with KRAS (**Fig. 3a**, lower panel). Calculation of the solvent-accessible area shows that 658 Å^2^ of RBD and 555 Å^2^ of CRD are buried when RBDCRD is in complex with KRAS, suggesting that both RBD and CRD have similarly sized interaction interfaces with KRAS. Sequence alignment of the linker region between RBD and CRD reveals that five of the six amino acid residues are conserved across all three RAF isoforms (**Fig. 3b**) suggesting similar interactions between RBD and CRD amongst all RAFs.

**Figure 3:**
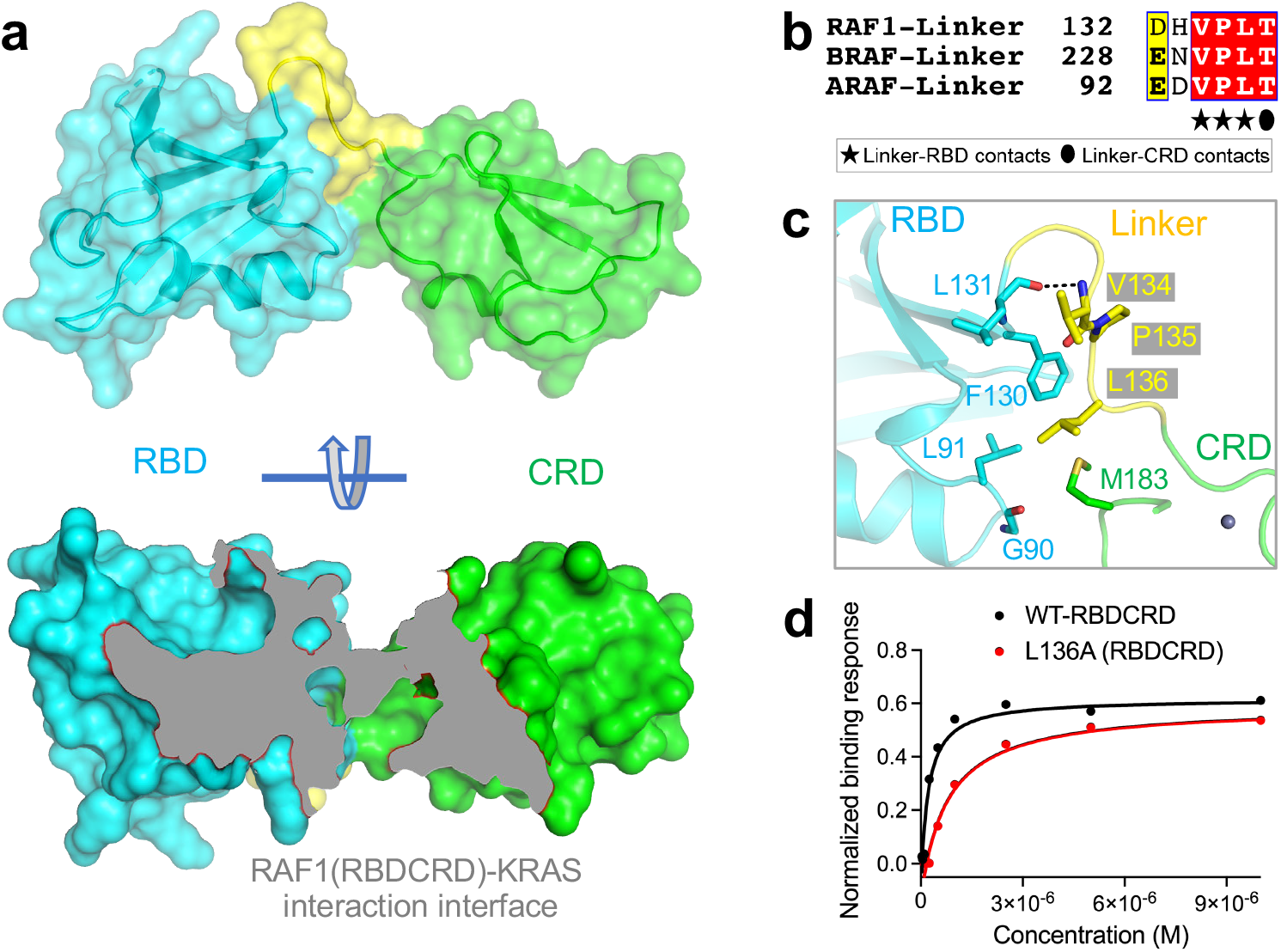
Structural and mutational analyses of the role of linker residues present between RBD and CRD in the KRAS-RBDCRD structure. **(a)** The surface representation of RBDCRD showing how linker residues connect RBD and CRD, resulting in the formation of an extended structure. The lower panel shows the region of RBDCRD that interacts with KRAS (colored gray), suggesting RBD as well as CRD interact with KRAS. **(b)** Amino acid sequence alignment of residues present in the linker region of human RAF1, BRAF, and ARAF. Residues that are involved in the interaction with RBD and CRD are indicated below the alignment with stars and oval, respectively. **(c)** Enlarged view of the RBD and CRD interaction interface formed by residues present in the linker region. Residues that participate in inter-domain interaction are shown in stick representation. A lone hydrogen bond is shown as a black dashed line. **(d)** Steady-state binding isotherms derived from the SPR data for the WT-RBDCRD and L136A mutant present in the linker region binding to KRAS-GMPPNP.

Interactions between the linker region and RBD involve a hydrogen bond and hydrophobic interactions between L131 of RBD and V134 of the linker region via atoms in the main chain and side chain, respectively (**Fig. 3c, Supplementary Table 4**). Furthermore, the phenyl ring of F130 (RBD) is present in a hydrophobic pocket formed by two linker residues P135 and L136. The linker and CRD interactions involve hydrogen bond and hydrophobic interactions by linker residue T137 and CRD residues T138, and C184 (part of the zinc-finger motif) present at the C-terminal end of CRD. In crystal form II, CRD interaction with the linker region also includes a hydrogen bond between residues P135 and D186. The structural superposition of the two crystal forms of the KRAS-RBDCRD complex shows conformational differences in the linker region (**Supplementary Fig. 1b**), partly due to crystal packing interactions but nonetheless shows that the short linker region plays a role in stabilizing RBD and CRD together as one extended structure. RBD and CRD also interact directly with each other via non-bonded interactions formed by residues G90 and L91 from RBD and M183 from CRD (**Supplementary Table 4**).

To examine the role of hydrophobic linker region residues in stabilizing the RBDCRD structure, we mutated residue L136 within the linker region to alanine. SPR analysis shows that this L136A mutation led to a 4-fold reduction in binding affinity to KRAS compared with WT-RBDCRD (**Fig. 3d, Supplementary Fig. 2d**). Considering that L136 is not present at the KRAS-RBDCRD interface, the 4-fold decreased binding affinity for this RBDCRD mutant with KRAS highlights the role of the linker residues in stabilizing RBDCRD and in facilitating interdomain interaction between RBD and CRD.

### RAF1(CRD) interacts with KRAS primarily via the inter-switch region and C-terminal helix α5

The overall structure of CRD within the tandem RBDCRD present in the crystal structures of the KRAS-RBDCRD complex resembles the previously reported NMR structure of the CRD domain^20^ (**Supplementary Fig. 3a**). The r.m.s. deviation between the crystallographic and NMR (minimized averaged, PDB 1FAR) structure is 1.6 Å for the Cα atoms. Secondary structure assignment for the CRD residues using the program DSSP showed three β-strands (β1-β3) forming an antiparallel β-sheet and a 3_10_-helix (η1) present in all structures described here. Similar to the solution structure^20^, our crystal structure contains two zinc finger motifs – one near the N- and C-terminal ends of CRD formed by residues H139, C165, C168, and C184, and a second one near the β3 strand and 3_10_-helix formed by residues C152, C155, H173, and C176 (**Supplementary Fig. 3b, c**).

The structures of the KRAS-RBDCRD complex provided, for the first-time, atomic details of the KRAS-CRD interaction interface, which is similar in size to the KRAS-RBD interface and also consists of eight hydrogen bonds. However, unlike the KRAS-RBD interface, which predominantly consists of polar and charged interactions, the KRAS-CRD interface contains no salt bridges and instead includes a relatively large hydrophobic interface (**Fig. 4a-c**). Interestingly, instead of the switch regions which have been shown to be involved in KRAS interactions with effectors, KRAS interacts with CRD mainly via residues present in the inter-switch region (R41, K42, Q43, V44, V45, I46, D47 and G48), and in the C-terminal helix α5 (R149, D153 and Y157) (**Supplementary Fig. 3d, e**). Eight of these eleven KRAS residues form hydrogen bonds with CRD residues (**Fig. 4d, e, Supplementary Table 5**). Although the switch regions of KRAS are not involved at the KRAS-CRD interface, KRAS residues L23, I24 and N26 present at the N-terminal end of the switch-I form non-bonded interactions at the KRAS-CRD interface. CRD residues that are involved at the KRAS-CRD interface are present at the N-terminal end of CRD (T138, H139, F141 and R143), β2-strand (F163), 3_10_-helix (E174, H175, S177, T178 and K179), and towards the C-terminal end of CRD (V180 and T182). Among these CRD residues, H139, R143, E174, S177, T178 and V180 form hydrogen bond interactions with KRAS. The hydrophobic amino acids in both KRAS and CRD interact with each other and enhance KRAS-RBDCRD interaction. The 3_10_-helix present in CRD is positioned parallel to the β2 strand and at the top of the α5 helix in KRAS, resulting in surface complementarity at the KRAS-CRD interface.

**Figure 4:**
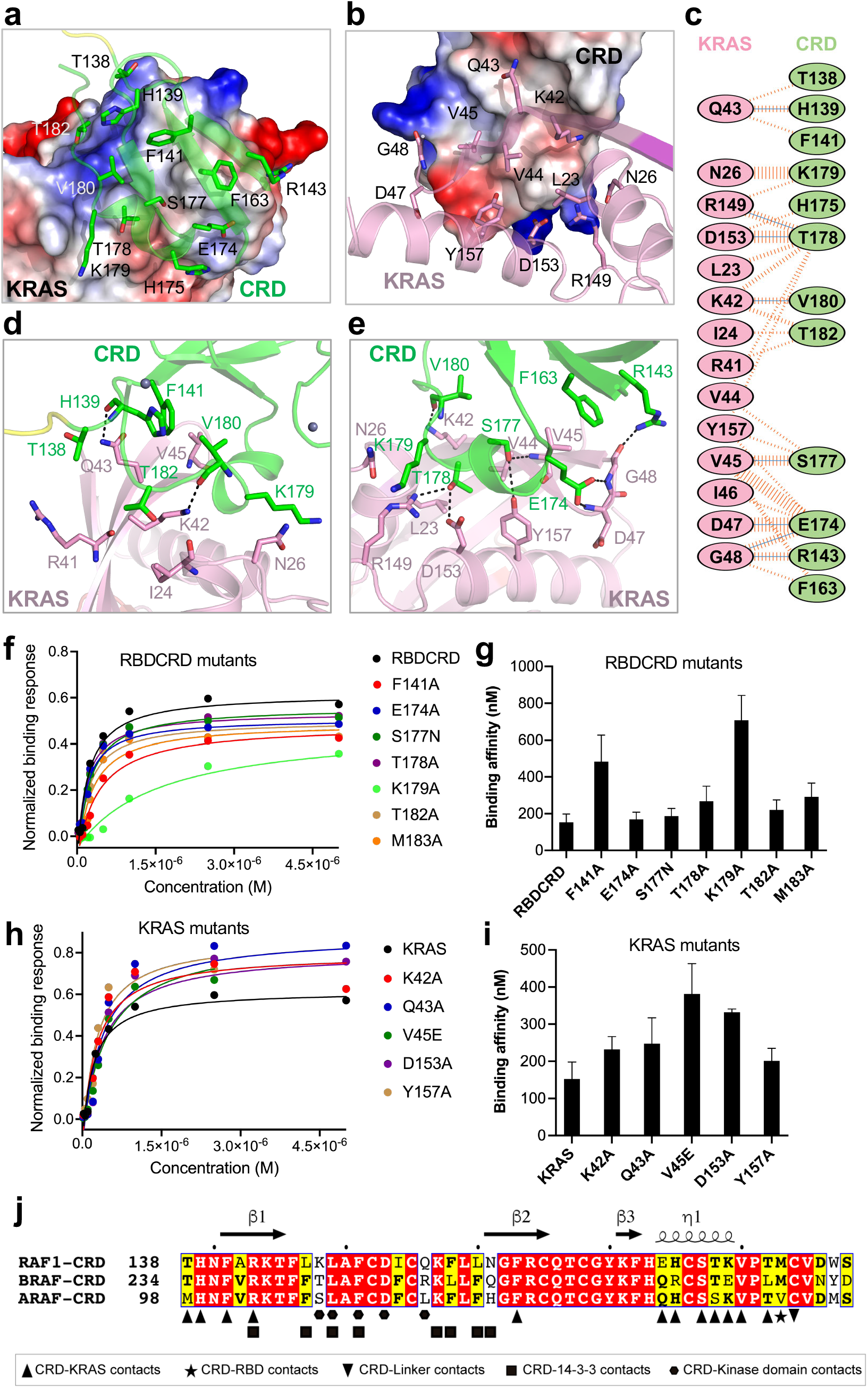
Structural and mutational analyses of the KRAS-CRD interactions in the KRAS-RBDCRD complex. **(a)** The KRAS-CRD interaction interface, with KRAS shown in electrostatic surface representation and the CRD residues that participate at the interface shown in stick representation. **(b)** The KRAS-CRD interaction interface, with CRD shown in electrostatic surface representation and the KRAS residues that participate at the interface shown in stick representation. **(c)** Schematic representation of the KRAS-CRD interaction interface, as identified by PDBSum. The interactions are colored using the following notation: hydrogen bonds – solid blue lines, salt bridge – solid red lines, non-bonded contacts – striped orange lines (width of the striped line is proportional to the number of atomic contacts). **(d, e)** Enlarged view of the KRAS-CRD interaction interface formed by residues present in the (d) inter-switch region, and (e) helix α5 of KRAS. The KRAS and CRD residues that participate in the protein-protein interaction are shown in stick representation. Intermolecular hydrogen bonds are indicated by dashed black lines. **(f)** Steady-state binding isotherms derived from the SPR binding kinetics for the point mutants of RBDCRD residues (present at the KRAS-CRD interface) binding to KRAS-GMPPNP. (**g**) A bar graph visualization of binding affinity (K_D_) obtained using the SPR data shown in (f) for point mutants of RBDCRD residues located at the KRAS-CRD interface. **(h)** Steady-state binding isotherms derived from the SPR binding kinetics for the mutants of KRAS residues (present at the KRAS-CRD) interface binding to WT-RBDCRD. **(g)** A bar graph showing binding affinity (K_D_) obtained using the SPR data shown in (h) for point mutants of KRAS residues located at the KRAS-CRD interface. (**j**) Amino acid sequence alignment of residues present in the cysteine-rich domain (CRD) of human RAF1, BRAF, and ARAF. Fully and partially conserved residues among the RAF isoforms are highlighted in red and yellow, respectively. The secondary structure of RAF1(CRD) is shown above the alignment. The RAF1(CRD) residues that are involved in the interaction with KRAS, RBD, and linker are indicated below the alignment with upright triangles, star and inverted triangles, respectively. In addition, based on the cryoEM structure of BRAF-MEK-14-3-3 (PDB 6NYB), CRD residues that are involved in the interaction with 14-3-3 and BRAF kinase domain are indicated with squares and hexagons, respectively.

To identify key residues that play a major role at KRAS-CRD interaction interface, we mutated hydrophobic and polar residues in both RBDCRD and KRAS, and measured their binding affinities using SPR. Among the CRD point mutants (F141A, E174A, S177N, T178A, K179A, T182A and M183A) that we tested, K179A and F141A showed 3-4-fold reduced affinity for active KRAS suggesting the importance of both hydrophobic and hydrogen bonding interactions at the KRAS-CRD interface (**Fig. 4f, g and Supplementary Fig. 4a**). All KRAS mutants (K42A, Q43A, V45E, D153A and Y157A) that we examined exhibited modest effects, with V45E and to a lesser extent D153A mutations resulting in ~2-fold reduction in the binding affinity (**Fig. 4h, I, and Supplementary Fig. 4b**). The relatively modest effect of mutations at the KRAS-CRD interaction interface suggests that, overall, the interactions between KRAS and RBD play a more significant role in the formation of KRAS-RBDCRD complex.

Amino acid sequence alignment of CRD residues in RAF isoforms shows significant sequence identity. Twelve out of fifty residues present in CRD are part of the KRAS-CRD interface (**Fig. 4j**). Among these twelve residues, six are fully conserved among all three RAF isoforms, whereas the remaining six are conserved among two of the three RAF isoforms suggesting a conserved RAS-CRD interface. In CRD, residue M183 interacts with RBD via hydrophobic interaction, while C184 interacts with T137 located in the linker region via hydrophobic interaction and hydrogen bonding with main chain atoms. Residue C184 is part of the zinc finger and is conserved among all RAF isoforms, whereas M183 is conserved in RAF1 and BRAF and replaced by another hydrophobic residue (valine) in the case of ARAF. A recently solved cryo-EM structure of autoinhibited BRAF in complex with 14-3-3 and MEK (PDB 6NYB) shows CRD occupying a central position in the complex by interacting with a 14-3-3 dimer and the kinase domain of BRAF^5^. Interestingly, except for R143, none of the 14-3-3- and kinase domain-interacting residues from CRD are part of the KRAS-CRD interface (**Fig. 4j**). This suggests that the KRAS-binding interface is likely to be partially exposed in the autoinhibited state of RAF.

### RAS-CRD interaction is important for RAS-mediated RAF activation

The point mutations in CRD and KRAS residues present at the KRAS-CRD interface do not significantly impact KRAS-RAF1 interaction, confirming previous reports that the KRAS-RBD interface plays a dominant role in the formation of the KRAS-RAF1 complex. To determine what role specific KRAS-CRD interactions play in RAS-mediated RAF activation, we examined the effects of point mutations in KRAS and RBDCRD by measuring KRAS-induced RAF1 kinase activity for these mutants. To measure the effects of mutations in RAF1, we co-expressed a series of point mutants with constitutively active KRAS Q61L. We purified each of them and measured their kinase activity at four concentrations, using MEK1 as a substrate (**Supplemental Fig. 5a**). **Figure 5a** shows that point mutations within CRD did not lead to substantial effects on binding to KRAS, consistent with SPR data described above. However, we observed a reduction (25-50%) in kinase activity with CRD or linker mutants, comparable to the effects of RBD mutants (**Fig. 5b**). Among the RAF1 mutants, T178A mutation located in CRD and L136A mutation located in the linker region led to the highest reduction (~50%) in kinase activity. Both residues potentially play important roles in proper CRD association with KRAS, with T178 forming two hydrogen bonds with KRAS (**Fig. 4c, e)** and L136 buried at the RBD-CRD interface (**Fig. 3c**), potentially orienting CRD to interact with KRAS. This suggests that in addition to the high affinity binding of RAF1 to KRAS provided by RBD, proper CRD association with KRAS is required for robust RAF1 activation.

**Figure 5:**
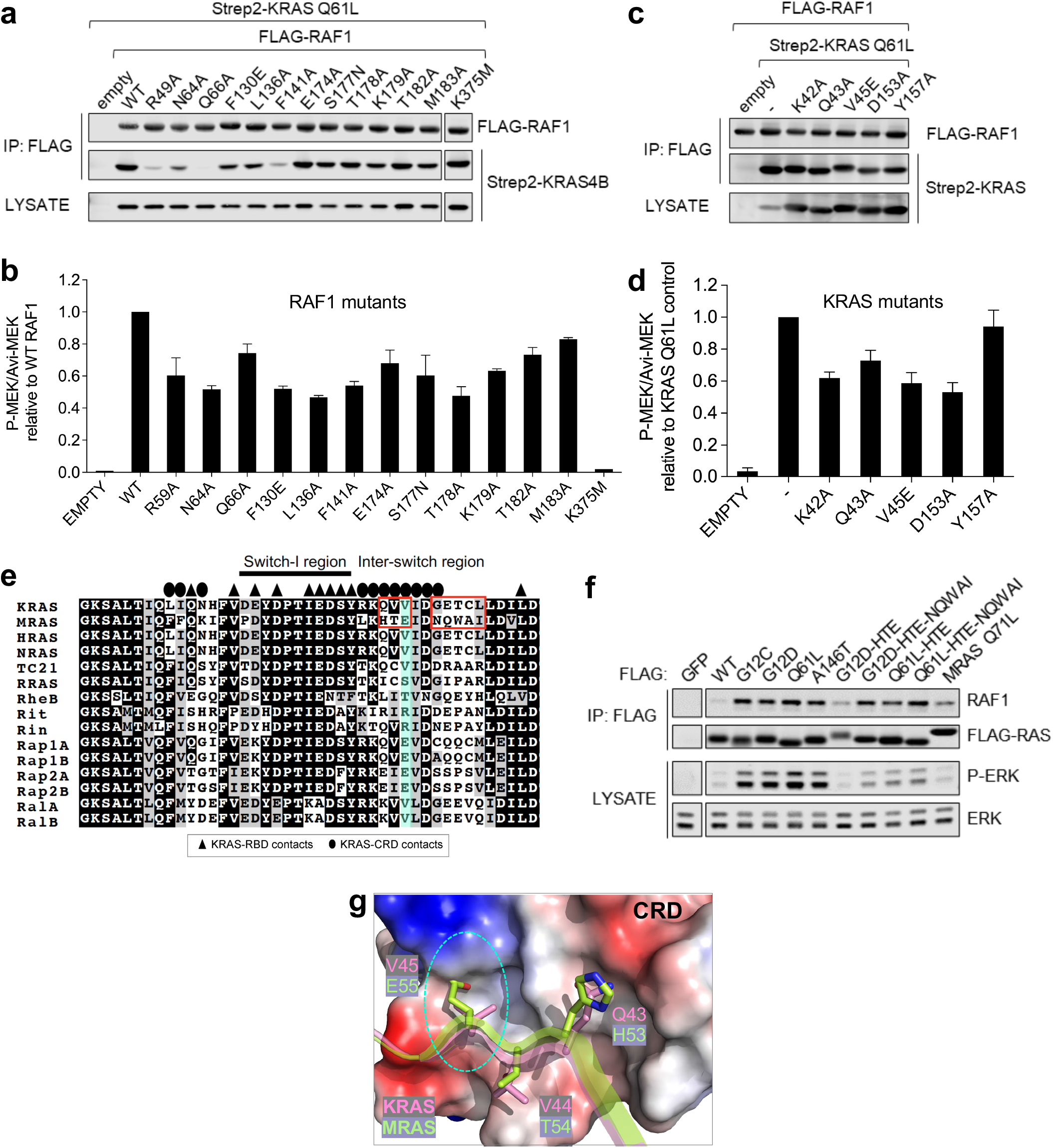
Interactions at the RAF1-CRD interface with KRAS are required for full RAF1 kinase activity. **(a)** FLAG-RAF1 WT and mutants for kinase assays were immunoprecipitated with FLAG antibody from 293T cells and assessed for binding to Strep2-KRAS. (**b**) Kinase activity of purified RAF1 mutants from cells co-expressing KRAS Q61L (shown in panel 5a). **(c)** FLAG-RAF1 WT and Strep2-KRAS mutants were co-expressed and isolated as in panel 5a. The “-” denotes KRAS-Q61L with no additional mutations. **(d)** Kinase activity of purified RAF1 co-expressed with KRAS mutants (shown in panel 5c). **(e)** Sequence alignment of residues present in the switch I, and inter-switch region of human RAS isoforms and members of RAS subfamily. Fully conserved residues are highlighted in black. The switch and inter-switch regions are indicated above the alignment. KRAS residues that are involved at the KRAS-CRD and KRAS-RBD interfaces are highlighted above the alignment using ovals and triangles, respectively. The red boxes and cyan color highlight a lack of conservation in the inter-switch region among RAS isoforms and MRAS as well as other members of the RAS subfamily. **(f)** FLAG-KRAS WT or mutant, or constitutively active MRAS-Q71L immune precipitates were probed for endogenous RAF1 interaction and pathway activation was measured in lysates. “HTE” denotes substitution of inter-switch residues ^43^QVV^45^ (KRAS) to HTE (MRAS), and “HTE-NQWAI” was changed from ^43^QVV^45^-^48^GETCL^52^ (KRAS) to HTE-NQWAI (MRAS). **(g)** The KRAS-CRD interaction interface, with CRD shown in electrostatic surface representation and the KRAS residues that participate at the interface shown in stick representation. Structural superposition of MRAS (PDB 1X1S) with KRAS shows residue E55 from MRAS clashes with CRD.

Similar experiments were performed to examine the ability of KRAS mutants to activate wild type RAF1. KRAS mutations were introduced into the constitutively active Q61L background and then co-expressed with RAF1 showed no significant changes in KRAS-RAF1 binding (**Fig. 5c**). With the exception of Y157A, all mutations were less efficient at activating RAF1 kinase (**Fig. 5d, Supplementary Fig. 5b**). These results suggest that besides the switch-I region in KRAS, residues present in the inter-switch region, especially residues K42 to V45, play an important role in RAS-mediated RAF activation, most likely through KRAS-CRD interactions.

Members of RAS subfamily GTPases such as RRAS, TC21, MRAS, Ral, RheB, Rap1, Rit and Rin share high sequence homology with three RAS isoforms in the switch-I region (**Fig. 5e**), the main interacting interface with RAF1(RBD). Despite this sequence homology, these RAS subfamily members show significantly reduced levels of RAF activation compared to RAS isoforms^26^. Interestingly, these members of the RAS subfamily diverge significantly in inter-switch regions with RAS isoforms (**Fig. 5e**). We hypothesize that the reduced RAF activation levels in other RAS family GTPases are due to the different residues in the inter-switch region, as this region accounts for the majority of the KRAS-CRD interactions. To test this hypothesis, we swapped residues from the inter-switch region in KRAS with that of MRAS, generating the HTE and HTE-NQWAI mutants in different constitutively active oncogenic KRAS backgrounds. While these mutants retain interaction with RAF1 as indicated in pull-down experiments (**Fig. 5f**), we observed a marked reduction in the ability of these variants to activate the ERK pathway, similar to that of constitutively active MRAS-Q71L. Structural alignment of MRAS (PDB: 1X1S) onto KRAS in the KRAS-RBDCRD complex suggests that among the inter-switch region mutations, V45E might have the highest impact on RAF1 activation, as the glutamate side chain will likely clash with CRD (**Fig. 5g**). The importance of V45 is further highlighted by our sequence alignment, which indicates that while V45 is conserved in RAS isoforms, over half of the other RAS subfamily GTPases have long chain amino acids (glutamate or arginine) instead of valine (**Fig. 5e**). Taken together, our results suggest that the inter-switch region of KRAS is important in fully activating RAF1 by interacting with CRD.

### Oncogenic mutants at G12, G13, and Q61 do not affect KRAS-RAF1(RBDCRD) interaction

To understand if oncogenic mutations at G12, G13, and Q61 positions in KRAS have any effect on KRAS-RBDCRD interaction interfaces, and to identify any druggable pockets at the KRAS-RBDCRD interface, we attempted to crystallize and solve the structures of G12C/D/V, G13D, and Q61L/R mutants of KRAS in complex with RAF1(RBDCRD). We were able to obtain well-diffracting crystals of G12V, G13D, and Q61R mutants of KRAS complexed with RAF1(RBDCRD) and solved their structures in the resolution range between 2.19-2.87 Å. (**Table 1, Supplementary Table 1**). Since the mutant structures belong to the same crystal form as the WT KRAS-RBDCRD structure obtained in crystal form II, we used this structure to examine any structural changes in KRAS and at KRAS-RBDCRD interfaces. Structural superposition of WT KRAS-RBDCRD with all three mutant KRAS-RBDCRD structures shows similar interaction interfaces, suggesting that the two RAS-binding domains of RAF1 bind to oncogenic KRAS mutants similarly to WT KRAS (**Fig. 6a**). A closer examination of the switch regions of KRAS shows minor conformational changes in the structures of oncogenic mutants complexed with RBDCRD when compared with WT KRAS-RBDCRD structure (**Fig. 6b**). The presence of a larger side chain at G12, G13, or Q61 position results in local rearrangement of side chains of some of the neighboring residues present in the switch regions without perturbing interactions at KRAS-RBDCRD interfaces. In the KRAS G12V-RBDCRD structure, KRAS residue Y32 adopts a different rotameric conformation to accommodate the side chain of Val at position 12. In the KRAS G13D-RBDCRD structure, the side chain of Y32 is disordered, suggesting multiple rotameric conformations of this residue to avoid a steric clash with the side chain of D13. The side chain of R61 in the KRAS Q61R-RBDCRD structure points towards the KRAS-RBD interface and occupies an empty space surrounded by KRAS residues Y64, P34, and I36, therefore not impacting the KRAS-RBD interface. These results suggest that RAF1 does not differentiate between WT and oncogenic mutants of active KRAS. Though there are no distinct new druggable pockets in oncogenic mutant structures with RBDCRD, these structures provide a molecular blueprint for designing peptidomimetics that are specific to individual oncogenic mutant allele of KRAS.

**Figure 6.**
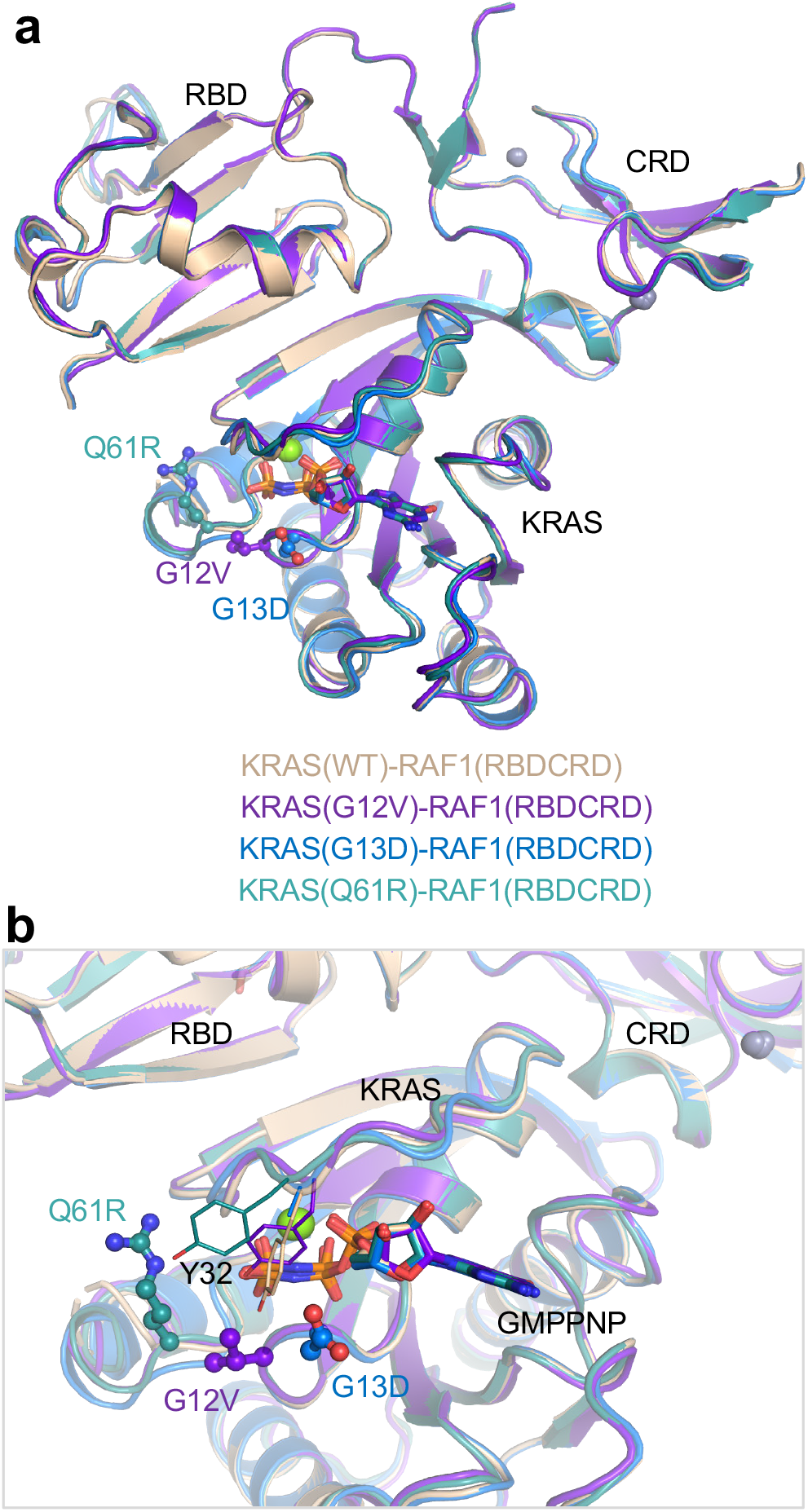
Structural comparison of WT KRAS-RBDCRD with the oncogenic mutants G12V, G13D, and Q61R of KRAS bound to RBDCRD. **(a)** Structural superposition of WT, G12V, G13D and Q61R mutants of KRAS complexed with RBDCRD. The structural superposition was carried out using KRAS residues. Crystal form II of WT KRAS-RBDCRD is used here since it has the same space group as the oncogenic mutants. Oncogenic mutants and GMPPNP are show in ball-and-stick and stick representations, respectively. The color scheme for different structures are shown in the panel. **(b)** Enlarged view showing oncogenic mutants, residue Y32 in the switch-I region, and KRAS-RBDCRD interface to highlight similarities and differences among structures of WT and oncogenic mutants of KRAS in complex with RBDCRD.

### Model showing the KRAS-RAF1(RBDCRD) complex at the membrane

Using ELISA, it was shown that RAF1(CRD) interacts selectively with phosphatidylserine (PS), and a cluster of basic amino acids, R143, K144, and K148 is critical for the CRD interaction with PS^7,27^. In addition, NMR titration and PRE experiments involving RAF1(CRD) binding to nanodisc showed chemical shift perturbations and PRE-induced peak broadening for basic residues K148 and K157, and hydrophobic residues around them^6,8^. Mapping these CRD residues in the KRAS-RBDCRD structure shows that these residues are fully-exposed on the CRD surface, allowing them to interact with the membrane while the CRD is bound to KRAS (**Fig. 7a**). The electrostatic surface representation of this region of CRD highlights a hydrophobic and basic patch on the surface that is likely to be involved in the interaction with PS-containing membranes (**Fig. 7b**).

**Figure 7.**
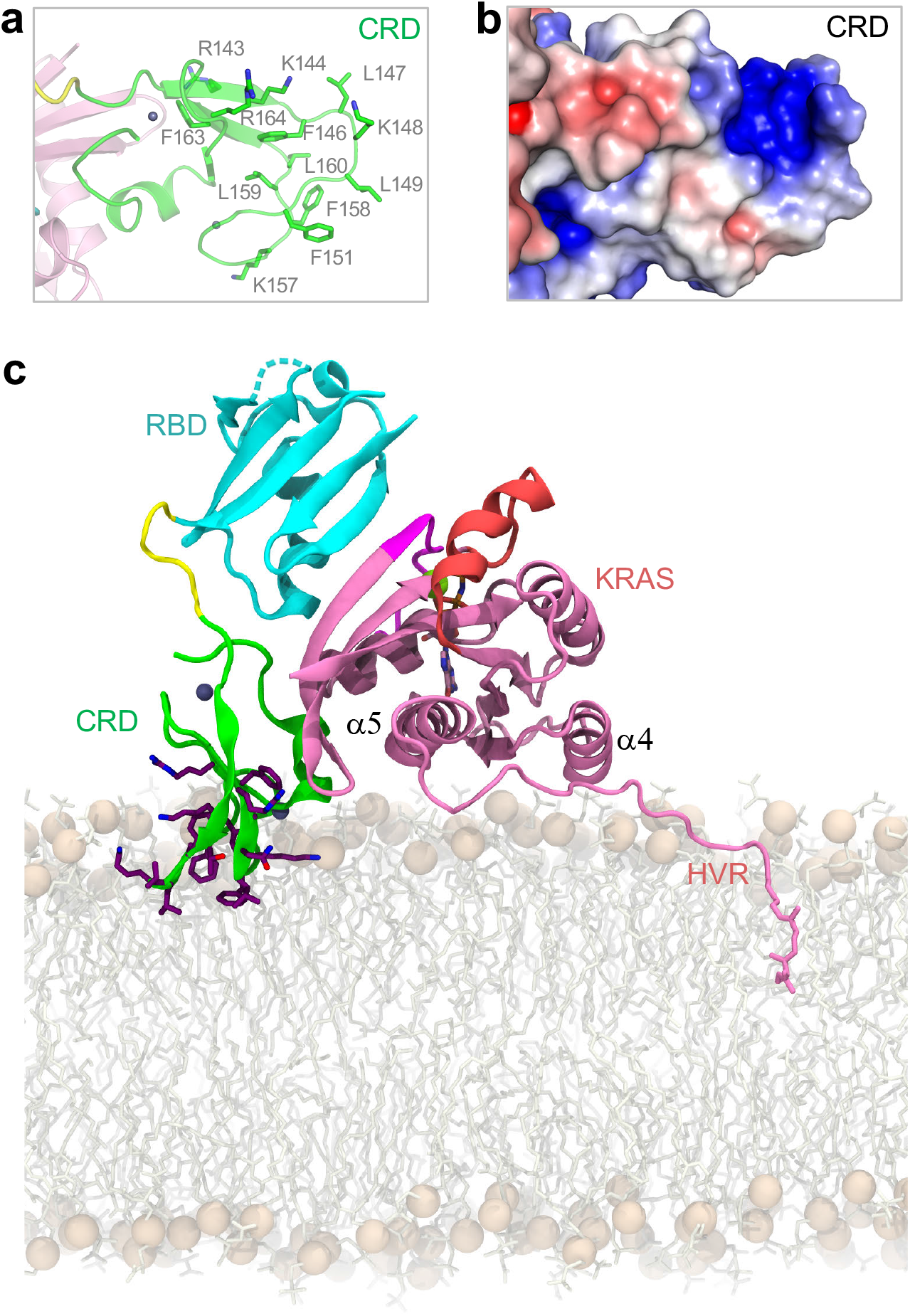
Membrane interacting residues in CRD, and a model showing the crystal structure of KRAS-RBDCRD at the membrane. **(a)** Enlarged view of the CRD part of the KRAS-RBDCRD structure with CRD residues (L147, K148, L149, A150, and F158) that have been shown to interact with the PS-containing membrane shown as sticks. **(b)** Electrostatic surface representation of the CRD shown in panel 7a highlighting a basic patch which has been proposed to interact with the PS headgroups on the membrane. **(c)** Crystallographic KRAS-RBDCRD oriented with KRAS helices α4 and α5 in membrane contact, as observed in MD simulations of KRAS^28^. Lipids are sticks with spheres for headgroup phosphorus atoms; KRAS-RBDCRD has the same color coding defined in Figure 1c. Side chains of CRD residues R143KTFLKLAF151 and K157FLLNGFR164 are sticks with purple carbon atoms.

To assess possible membrane interactions of our KRAS-RBDCRD structure, we constructed several membrane-bound active KRAS-RBDCRD models. Previously, MD simulations have identified three highly populated orientations of KRAS on a membrane containing 30% anionic PS lipids in which the region of RAS that approaches the cell membrane is described by a rotation angle^28^ (**Supplementary Fig. 6a-c**). We aligned our crystal structure of KRAS-RBDCRD to these three KRAS poses and evaluated them based on agreement with CRD residues that have been shown to interact with the membrane^8^. Specifically, the relative arrangement of protein domains in our crystal structure indicates that membrane interaction of KRAS helices α4 and α5 (rotation angle=-15°) is associated with membrane insertion of two CRD loops composed primarily of hydrophobic and cationic residues (**Supplementary Fig. 6d, e**), in agreement with NMR data^8,29^ (**Fig. 7c).** In contrast, while membrane approach of KRAS β-strands β1-β3 (rotation angle=105°) allows concerted membrane interaction of KRAS, RBD, and CRD, these orientations exist on the brink of membrane occlusion (**Supplementary Fig. 6f**) involving RBD and linker regions of RAF (**Supplementary Fig. 6g**), and are incompatible with membrane insertion of the CRD’s hydrophobic loops in the absence of inter-domain reorganization (**Supplementary Fig. 6g**). Furthermore, KRAS orientation such that helices α3 and α4 approach the membrane (rotation angle=-85°) appears incompatible with existing experimental data since this orientation of KRAS places CRD far from the membrane (**Supplementary Fig. 6h**). We note that our model of membrane-bound KRAS-RBDCRD shown in **Fig. 7c** is similar to the State A configuration proposed recently by Fang *et al*.^29^ based on an NMR data-driven model generated using docking software HADDOCK, excepting a 180° rotation of CRD about the approximate RBDCRD long axis and slightly more membrane insertion of CRD loops due to greater engagement of KRAS helix α5 with the membrane surface (**Supplementary Fig. 6i**). This discrepancy of the CRD orientation could be due to the presence of a nanodisc in the NMR model, insufficient restraints on the CRD’s β1 and 3_10_-helix (since many residues in these regions were broadened beyond detection), and/or perhaps higher flexibility of CRD in solution in general as indicated by the many broadened peaks. The membrane-bound KRAS-RBDCRD model shown in Fig. 7c represents a combination of our high-resolution crystal structure with a preferred membrane orientation of KRAS according to MD simulations and NMR data. This model also positions previously identified CRD membrane-interacting residues into the lipid bilayer and may therefore provide a snapshot of RAF1 activation by KRAS at the membrane.

## DISCUSSION

The RAS-RAF interaction has taken center stage in cancer research as it is implicated in almost 20% of all known human cancers^30^. Direct protein-protein interaction between RAS and the N-terminal regulatory domain of RAF is critical for the recruitment of RAF to the plasma membrane, a crucial step in the process of RAS-mediated RAF activation^1,2^. Even though the RAS-RAF interaction was discovered more than 25 years ago, it remains unclear how RAS interacts with RAF during the multi-step process of RAF activation. Recent reports of cryoEM and crystal structures of the full-length BRAF or BRAF kinase domain in complex with 14-3-3 and/or MEK have provided new insights into the autoinhibited and active states of RAF^5,22,31,32^. However, in these structures, the entire N-terminal half of BRAF is either not observed or, in the case of the autoinhibited complex, only the CRD can be seen centrally anchored between 14-3-3 and the kinase domain of BRAF^5^. Moreover, these structures do not contain RAS and therefore provide limited insights into RAS-RAF interaction during the process of RAS-mediated RAF activation.

The crystal structure of KRAS in complex with RBDCRD of RAF1 described here shows how the structure of tandem domains RBD and CRD form one structural entity, and how the two RAS binding domains of RAF interact with KRAS. This structure reveals that the KRAS-RBD binding interface is similar to that of HRAS-RBD and Rap1-RBD structures^19,21^, as well as KRAS-RBD in the absence of CRD. Importantly, our KRAS-RBDCRD structure provides atomic details of the KRAS-CRD interaction interface, which supports the previous mutagenesis experiments where KRAS residues N26 and V45 were suggested to contribute to RAS-RAF interaction and RAF activation^12^. Previous mutagenesis studies on RAF1(CRD) have also shown the complete loss of RAS-mediated RAF activation in the CRD double mutants K144A and L160A^9^. These two CRD residues are located in the hydrophobic loops, where they play an important role in anchoring CRD to the membrane. Using the KRAS-RBDCRD structure and MD simulations of KRAS on a membrane with 30% PS^28^, we constructed a membrane-bound KRAS-RBDCRD model that has K144, L160 and other residues in the CRD hydrophobic loops inserted into the membrane.

The crystal structure of the KRAS-RBDCRD complex and structural insights obtained by comparing it with the autoinhibited structure of BRAF in complex with 14-3-3/MEK provide new insights into the process of RAS-mediated RAF activation (**Fig 8a-c**). In the autoinhibited state, BRAF and MEK have been shown to exist as a preassembled quiescent complex bound to 14-3-3^33^. Recent cryoEM structures by Park *et al*. showed that the BRAF-MEK1 complex is secured in a cradle formed by the 14-3-3 dimer, which binds the two phosphorylated serine sites flanking the BRAF kinase domain^5^. The CRD of BRAF occupies a central position where its membrane-binding surface is buried by interactions with the 14-3-3 dimer and the kinase domain. The molecular handcuffing of BRAF by 14-3-3 therefore maintains the autoinhibited state by blocking the kinase domain’s dimerization and preventing CRD from binding to the membrane. Interestingly, structural superposition of our KRAS-RBDCRD structure to the autoinhibited complex via CRD residues places RBD near the residual electron density observed in the autoinhibited complex, which Park *et al*. suspected to represent the RBD^5^ (**Fig. 8a**). In this superposed structure, KRAS is able to bind to the exposed RBD without sterically clashing with neighboring 14-3-3 or kinase domain residues (**Fig. 8b**).

**Figure 8.**
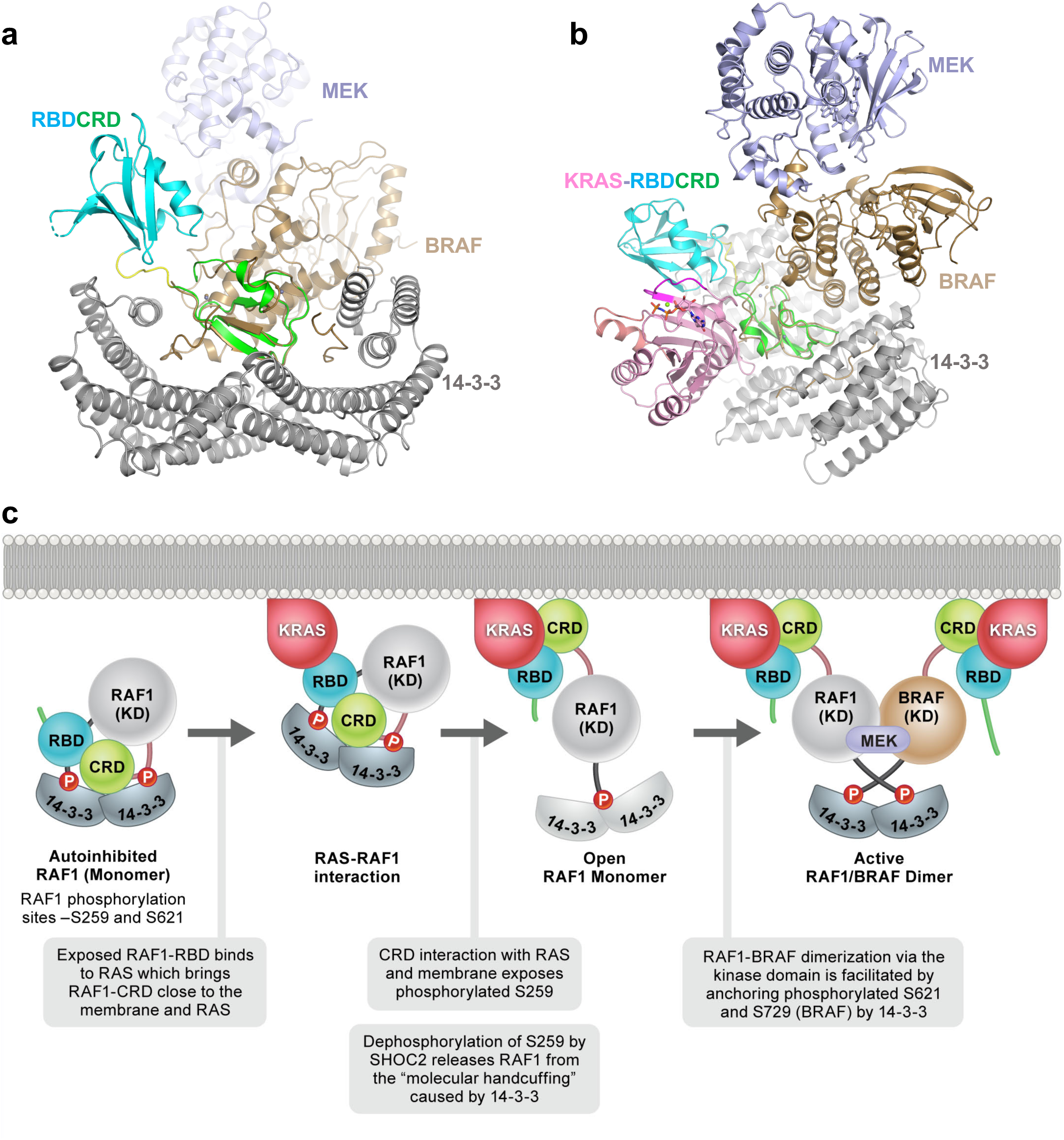
Model of RAS-mediated RAF activation using structural insights obtained from KRAS-RBDCRD and MEK-BRAF-14-3-3 structures. **(a)** Structural superposition of KRAS-RBDCRD and the cryoEM structure of MEK-BRAF-14-3-3 (PDB 6NYB). Structures were aligned by their CRD domains. KRAS is hidden in this panel in order to show the alignment of CRD and the placement of RBD clearly. RAF1(RBDCRD) has the same color-coding scheme as used in Figure 1c, while MEK, BRAF, and 14-3-3 are colored light blue, brown, and gray, respectively. **(b)** Same as (a) but at a different angle. KRAS is shown to illustrate it has minimal contacts with 14-3-3 in our structural alignment based on CRD. **(c)** Schematic diagram showing KRAS first interacts with autoinhibited RAF via RBD and then CRD to bring RAF close to the membrane. This results in CRD coming out of the autoinhibited conformation and interacting with the membrane and KRAS, and in turn exposing the phosphorylated serine present in the CR2 region. Dephosphorylation of S259 (RAF1) by SHOC2 complex results in an active RAF1 monomer, where 14-3-3 binds to only one phosphorylated serine located at the C-terminal end. Homo- or hetero-dimerization of KRAS-RAF1 with an active RAF monomer containing either RAF1 or BRAF results in the formation of an active dimerized RAF complex. RAF1 has been shown to prefer hetero-dimerization with BRAF which is bound to MEK for initiating the activation cascade.

Almost all of the CRD residues that interact with KRAS are also exposed in the autoinhibited state and could interact with KRAS. However, the loop containing the phosphorylated serine (S621) at the C-terminus of the kinase domain partially clashes with the KRAS-CRD interaction. In our model of RAF activation by RAS, the activation process starts with the interaction between active KRAS (GTP-bound) and RBD which is exposed in the autoinhibited RAF complex (**Fig. 8c**). The KRAS-RBD interaction is followed by CRD’s release from the autoinhibited complex and its interaction with KRAS and the plasma membrane. It has been suggested that CRD extraction upon RAS binding and its localization to the membrane is the critical event that triggers the release from the molecular handcuffing caused by the 14-3-3 dimer^5^. The release of CRD exposes phosphorylated serine (S259) in the central CR2 region of RAF to be dephosphorylated by the SHOC2-PP1C complex^34^, resulting in the formation of an active RAF monomer, where 14-3-3 can no longer bind to CR2 but presumably still binds to the phosphorylated serine in the C-terminal region of RAF.

As shown in the cryoEM structures of the active BRAF complex, 14-3-3 now binds to C-terminal phosphorylated S729 (equivalent to S621 in RAF1) coming from two BRAFs, thus facilitating the formation of an active BRAF dimer. This is supported by recent crystal structures of the BRAF kinase domain in complex with 14-3-3, which showed that dimeric 14-3-3 induces dimerization of the BRAF kinase domain and increases BRAF activity by relieving the negative regulatory effect of ATP which prevents BRAF dimerization^32^. Unlike BRAF, which tends to form homodimers, RAF1 has been shown to prefer to heterodimerize with BRAF^35–37^. Therefore, the formation of the active RAF1 complex is likely to involve heterodimerization between monomeric RAF1 and monomeric BRAF bound to MEK.

Our work, and the work of others, has revealed that CRD plays a crucial role in stabilizing both the autoinhibited and active states of RAF. The importance of this domain is also indicated by the observation that more than 40% of activating BRAF mutations identified in RASopathy syndromes occur in the CRD^38^. Despite being only 50 residues long, CRD shows enormous versatility in that it can interact with the membrane, RAS, 14-3-3, and the RBD and kinase domain of RAF. Discrete interfaces on CRD are arranged in a way that allows this domain to stabilize the autoinhibited state by burying membrane-interacting residues, or conversely facilitate the active state by presenting membrane anchoring residues. Through these interactions, CRD plays a central role in the process of RAS-dependent RAF activation. This role is distinct from its role in binding to RAS, as suggested previously by mutagenesis and biochemical analysis. Mutations in CRD (S177, T182) that prevent interaction with the inter-switch region of RAS were described that retain RAS binding via RBD but are defective in kinase activation or transforming ability^9^. Here, we extend these early observations to show that multiple mutations in the inter-switch region of RAS or in CRD can result in impaired kinase activation, often with minimal effects on binding. While RAS has been described as a binary switch, our characterization of CRD indicates that it also plays a binary role in the activation-inactivation cycle of RAF kinases.

The distinction between binding and activation may explain why some members of the RAS superfamily fail to fully activate RAF kinases, despite being able to bind to RAF through the highly conserved switch-I region. We show that, for example, failure of CRD to engage the interswitch region of MRAS may account for MRAS’ failure to activate RAF efficiently. We speculate that the unique inter-switch regions of RAS family members may provide specificity for different effector functions, despite the shared switch-I sequences.

Efforts to disrupt binding of RAS to RAF(RBD) as a therapeutic strategy have, so far, been unsuccessful. The high-affinity RAS-RBD interface, which consists of anti-parallel β-strands, does not offer any obvious pockets to which a small molecule could bind with high affinity. It is also reasonable to expect nonspecific consequences when targeting the RAS-RBD interaction given that this this interface is conserved amongst members of the RAS superfamily. Conversely, targeting the interaction between CRD and RAS, or CRD and the plasma membrane, may offer novel therapeutic opportunities, considering their lower binding affinities, the specificity afforded by the inter-switch region, and the more flexible nature of the protein interfaces.

## METHODS

### Cloning, expression, and purification of recombinant proteins

#### Cloning

Constructs for protein expression were produced using Gateway recombination-based cloning as described previously^39^ using attB-flanked *E. coli* optimized synthetic DNA generated by ATUM, Inc. as starting materials. RAF1(RBDCRD) clones consisted of RAF1 amino acids 52-188 (or 52-192) with an upstream TEV protease site (ENLYFQ); cleavage of this protein with TEV protease leaves the native serine at amino acid 52 as the N-terminal amino acid of the final protein. RAF1(RBD) clone consisted of RAF1 amino acids 52-131 with an upstream TEV protease site (ENLYFQ); cleavage of this protein with TEV protease leaves the native serine at amino acid 52 as the N-terminal amino acid of the final protein. Wild-type and mutants of KRAS4b(1-169) clones were generated with an upstream tobacco etch virus (TEV) protease site (ENLYFQG), which when cleaved during purification leaves an additional glycine residue at the N-terminus of the KRAS4b protein. Avi-tagged clones were generated with an upstream TEV site (ENLYFQ) followed by the Avi tag sequence (GLNDIFEAQKIEWHEG) and the KRAS4b(1-169) sequence. In all cases, final protein expression constructs were generated by Gateway LR recombination into pDest-566 (Addgene #11517), an *Escherichia coli* T7–based expression vector based on pET42 and incorporating hexa-histidine (His6) and maltose-binding protein (MBP) tags at the N-terminus of the protein. Inactive human MEK1 (MAP2K1) with K97R mutation was synthesized by ATUM as an *E. coli* optimized Gateway Entry clone containing an upstream strong *E. coli* ribosome-binding site, and a downstream Avi-TEV-His6 fusion tag GLNDIFEAQKIEWHEGENLYFQGHHHHHH). Expression constructs were generated by Gateway LR recombination into pDest-521, a pET21 variant vector with a T7 promoter, and no fusion tags.

#### Protein expression

All RAF1(RBDCRD) proteins were expressed using the protocols outlined previously for RAF1 (52-188)^40^. KRAS, Avi-KRAS, RAF1(RBD), and MAP2K1 K97R-Avi-TEV-His6 proteins were expressed following protocols described in Taylor *et al*. (Dynamite media protocol, 16°C induction)^41^. Essentially, an overnight 37°C culture (non-inducing MDAG-135 medium) of the *E. coli* strain harboring the expression plasmid of interest was used as seed culture to inoculate (2% v/v) expression-scale cultures of Dynamite medium. The expression culture was grown at 37°C until OD600 reached 6-8, protein expression was induced with 0.5 mM IPTG, the culture was incubated at 16°C for 18-20 h, and the cells were harvested by centrifugation.

#### Protein purification

All RAF1(RBDCRD) proteins were purified following the protocols described in Lakshman *et al*.^40^, whereas KRAS, RAF1(52-131), and MAP2K1 K97R-Avi-TEV-His6 proteins were purified as outlined for KRAS4b(1-169) in Kopra *et al*^42^. Briefly, the expressed proteins of the form His6-MBP-TEV-target were purified from clarified lysates by IMAC, treated with His6-TEV protease to release the target protein, and the target protein separated from other components of the TEV protease reaction by the second round of IMAC. Proteins were further purified by gel-filtration chromatography in a buffer containing 20 mM HEPES, pH 7.3, 150 mM NaCl, 2 mM MgCl2 (GTPases only), and 1 mM TCEP. The peak fractions containing pure protein were pooled, flash-frozen in liquid nitrogen, and stored at −80°C.

### Crystallization and data collection

Purified GDP-bound KRAS proteins were exchanged to GMPPNP-bound forms using the procedure reported earlier^43^. Wild type and oncogenic mutants (G12V, G13D, and Q61R) of KRAS were used for the crystallization experiments. The GMPPNP-bound KRAS proteins were mixed with RAF1(RBDCRD) or RAF1(RBD) in a 1:1.2 stoichiometric ratio and incubated for approximately 30-60 min. The protein-protein complex was then passed through a size-exclusion column (Superdex75, 26/60, GE Healthcare) to remove unbound proteins. Crystallization screening was carried out at 20°C using the sitting-drop vapor diffusion method by mixing purified protein-protein complex (~9 mg/ml) with an equal volume of reservoir solution. Crystallization hits obtained from initial screening were optimized to improve the diffraction quality by systematically varying the pH, individual component concentrations, and the presence of additive and detergents (**Supplementary Table 1**). KRAS constructs with C118S mutation was also used to improve the resolution during screening and optimization as this mutation presumably reduces inadvertent cysteine oxidation during crystallization. Optimized crystals were harvested for data collection and cryoprotected with 25% (v/v) glycerol solution before being flash-frozen in liquid nitrogen. Diffraction data sets were collected at 24-ID-C/E beamlines at the Advanced Photon Source (APS), Argonne National Laboratory. The crystals of KRAS and KRAS-C118S complexed with RAF1(RBDCRD) belong to different space groups and diffracted to a resolution of 2.5 and 2.15 Å, respectively. Crystals of WT KRAS with RAF1(RBD) diffracted to a resolution of 1.4 Å, and oncogenic mutants of KRAS with RAF1(RBDCRD) diffracted to a resolution of 2.19-2.87 Å. Crystallographic datasets were integrated and scaled using XDS^44^. Crystal parameters and data collection statistics are summarized in **Table 1**.

### Structure determination and analysis

The structure of WT KRAS in complex with RAF1(RBD) was solved by molecular replacement using the program Phaser as implemented in the Phenix suite of programs^45,46^, with a protein-only structure of HRAS-RAF1(RBD) complex^19^ as a search model (PDB ID: 4G0N). The structures of WT KRAS in complex with RAF1(RBDCRD) in crystal form I (2.15 Å) and II (2.5 Å) were solved using the protein-only structure of KRAS-RAF1(RBD) complex and the NMR structure of RAF1(CRD) (PDB ID: 1FAR)^20^ as search models. The structures of oncogenic mutants (G12V, G13D, and Q61R) of KRAS in complex with RAF1(RBDCRD) were solved using the 2.5 Å WT KRAS-RAF1(RBDCRD) structure as the search model. The initial solution obtained from molecular replacement was refined using the program Phenix.refine within the Phenix suite of programs^45^, and the resulting *Fo-Fc* map showed clear electron densities for the GMPPNP nucleotide, KRAS, and RBD/RBDCRD domains. The model was further improved using iterative cycles of manual model building in COOT^47^ and refinement with Phenix.refine. GMPPNP nucleotide was placed in the nucleotide-binding pocket and followed by the addition of solvent molecules by the automatic water-picking algorithm in COOT. These solvent molecules were manually checked during model building until the final round of refinement was completed. Refinement statistics for the structures are summarized in **Table 1**. All of the structural figures were rendered in PyMOL (Schrödinger, LLC) or VMD^48^ with secondary structural elements assigned using the DSSP server (http://swift.cmbi.ru.nl/gv/dssp/). The amino acid sequence alignments were carried out using Clustal Omega^49^, and the figures were produced using ESPript^50^. Crystallographic and structural analysis software support is provided by the SBGrid consortium^51^.

### SPR measurements

SPR binding experiments were collected on a Biacore T200 or S200 Instrument (GE Healthcare). Neutravidin (Pierce) was amine coupled to the carboxymethylated dextran surface of a CM5 sensor chip (GE Healthcare) using standard amine coupling chemistry. The CM5 chip surface was first activated with 0.1 M N-hydroxysuccinimide and 0.4 M N-ethyl-N’-(3-dimethylaminopropyl) carbodiimide at a flow rate of 20 μl/min using 20 mM HEPES pH 7.4, 150 mM NaCl as the running buffer. Next, neutravidin was diluted to 20 μg/ml in 10 mM sodium acetate (pH 4.5) and injected on all 4 flow cells until a density of approximately 10,000 Response Units (RU) was immobilized. Activated amine groups were quenched with an injection of 1 M ethanolamine (pH 8.0). 100-400 RU of WT or mutant Avi-tagged KRAS (1-169) was captured on three flow cells in 20 mM HEPES pH 7.4, 150 mM NaCl, 5 mM MgCl2, 1 mM TCEP, 5 μM GMPPNP, 0.01% tween 20 buffer. RAF1 RBD (52-131), RBDCRD (52-192) and RBDCRD mutant proteins were diluted from 20 – 0.05 μM in 20 mM HEPES pH 7.4, 150 mM NaCl, 5 mM MgCl2, 1 mM TCEP, 5 μM GMPPNP, 0.01% tween 20 buffer and injected over all of the flow cells at 30 μl/min. Flow cell 1 was used for referencing, and buffer injections were included for referencing purposes. SPR sensorgrams were normalized by the capture level of KRAS to allow direct comparison between different experimental runs. Steady-state levels of RBD or RBDCRD were recorded and fit with a 1:1 binding model using the Biacore Evaluation software.

### Immunoprecipitation/pulldown and kinase assays

FLAG-RAF1 and Strep2-KRAS4b or empty vector mammalian expression plasmids were transiently expressed in 293T using JetPRIME transfection reagent according to the manufacturer protocol. Thirty-six hours later cells were washed with PBS and lysed on ice in 25 mM Tris pH 7.5, 150 mM NaCl, 1% Triton X-100, 5 mM MgCl2, 1 mM DTT, EDTA-free protease inhibitors (Roche) and phosphatase inhibitor cocktail (Millipore-Sigma). After centrifugation to clear the lysates, anti-FLAG M2 (Millipore-Sigma) or StrepTactin agarose (Novagen) were used to isolate tagged-protein complexes by rotating for 2 h at 4°C. Beads were washed with 25 mM Tris pH 7.5, 150 mM NaCl, 1% Triton X-100, 5 mM MgCl2. A portion of the beads was analyzed by SDS-PAGE and Western blot, where complexes were visualized using antibodies to phosphorylated, tagged or endogenous proteins, and DyLight-conjugated secondary antibodies suitable for Li-COR Odyssey scanning. The remaining FLAG beads were incubated with 150 ng/μL 3X FLAG peptide (Millipore-Sigma) in kinase assay buffer (100 mM Tris pH 7.5, 150 mM NaCl, 5 mM MgCl2, 1 mM DTT) for 10 min with agitation at 22°C to elute proteins. The supernatant was collected, and the elution was repeated twice, followed by one wash with buffer without peptide to remove traces of eluate from the beads. To perform the kinase assay, FLAG-RAF1 eluate was serially diluted with kinase assay buffer and combined with recombinant MEK1-K97R-Avi (final [10 ng/μL]) and 1 mM ATP. Reactions were agitated at 22°C for 20 min and were stopped with the addition of 4X NuPAGE LDS (ThermoFisher). A fraction of each assay sample was analyzed by Western blot as above. Quantification of band intensity was performed with ImageStudio Lite (Li-COR) software, and error bars represent S.D. of 3 independent experiments.

### Antibodies

Avi-tag - GenScript A01738; BRAF - SCBT sc-5284; ERK - Cell Signaling 9107; ERK P-Y202/Y204 - Cell Signaling 4370; FLAG - Sigma F7425; MEK P-S217/221 - Cell Signaling 9121; RAF1 - BD Transduction Laboratories 610152; NWSHPQFEK (Strep-tag) - GenScript A00626.

### Modeling KRAS-RBDCRD complex at the membrane

The KRAS-RBDCRD crystal structure was aligned such that the position and orientation of the KRAS G-domain matched configurations commonly observed in all-atom MD simulations of KRAS on 7:3 POPC:POPS bilayers^28^. Quantitative definition of KRAS orientation is outlined in **Supplementary Fig. 6a** and **6b**, and common orientations from simulation and experiment are shown in **Supplementary Fig. 6c**. Representative KRAS orientations from simulation were selected to maximize their similarity to the orientational centroid of that cluster and the average orientation-specific (± 5° tilt and rotation) displacement of the KRAS G-domain from the center of mass of the lipid bilayer along its normal, *d*_Gz_. KRAS fitting involves Cα atoms of KRAS G-domain residues Y4-N26, Y40-L56, and D69-H166 (selected to omit the relatively flexible N-terminus and switch regions of KRAS).

### Accession Numbers

The atomic coordinates and structure factors of the various complexes have been deposited in the Protein Data Bank and are available under accession numbers: 6XI7 – KRAS in complex with RAF1(RBDCRD), crystal form I; 6XHB – KRAS in complex with RAF1(RBDCRD), crystal form II; 6VJJ – KRAS in complex with RAF1(RBD); 6XHA – KRAS-G12V in complex with RAF1(RBDCRD); 6XGV – KRAS-G13D in complex with RAF1(RBDCRD), and 6XGU-KRAS-Q61R in complex with RAF1(RBDCRD).

## ACKNOWLEDGMENT

We thank Bill Gillette, Jennifer Mehalko, Rosemilia Reyes, Vanessa Wall, Jose Sanchez Hernandez, Nitya Ramakrishnan, Allison Champagne, Peter Frank, Min Hong, Taylor Lohneis, Shelley Perkins, Stephanie Widmeyer, Matt Drew, Peter Frank, and Kelly Snead of the Protein Expression Laboratory (Frederick National Laboratory for Cancer Research) for their help in cloning, expressing, and purifying recombinant proteins. We are grateful to the staff of 24-ID-C/E beamline at the Advanced Photon Source, Argonne National Laboratory, for their help with data collection. Part of this work is based on research conducted at the Northeastern Collaborative Access Team beamlines, which are funded by the National Institute of General Medical Sciences from National Institutes of Health Grant (P30 GM124165). The Eiger 16M detector on 24-ID-E is funded by a NIH-ORIP HEI grant (S10OD021527). This research used resources of the Advanced Photon Source, a US Department of Energy (DOE) Office of Science User Facility operated for the DOE Office of Science by Argonne National Laboratory under Contract DE-AC02-06CH11357. This project was funded in part with federal funds from the National Cancer Institute, National Institutes of Health Contract HHSN261200800001E. The content of this publication does not necessarily reflect the views or policies of the Department of Health and Human Services, and the mention of trade names, commercial products, or organizations does not imply endorsement by the US Government.

## Author contribution

T.H.T., A.H.C., and S.D. carried out structural work under the supervision of D.K.S.; L.C.Y. performed the immunoprecipitation/pulldown and kinase assays under the supervision of F.M.; L.B. carried out SPR experiments under the supervision of A.G.S.; and S.M., T.T., J-. P.D., and D.E. prepared clones and recombinant proteins. C.N. carried out modeling studies. D.K.S., D.V.N., and F.M. provided resources and contributed to the analysis. T.H.T., A.H.C., F.M., and D.K.S. wrote the manuscript with inputs from all co-authors. D.K.S. supervised and coordinated the overall project.

## Supplementary Information

### SUPPLEMENTARY METHODS

#### Quantification of RAS orientation

The orientation of KRAS with respect to the membrane surface is defined by two angles: the tilt angle, which separates the long axis of helix α5 from the bilayer normal, and the rotation angle, which defines which parts of KRAS are brought toward the membrane via tilting^8,28^ (**Supplementary Fig. 6a, b**). At tilt=0°, KRAS helix α5 is perpendicular to the membrane and all rotation angles are equivalent. At tilt=90°, KRAS helix α5 is parallel to the membrane surface and angles of rotation=-15°, −85°, and 105° dispose KRAS helices α4/α5, helices α3/α4, and β strands β1-3 toward the membrane, respectively (**Supplementary Fig. 6b**).

#### Membrane occlusion

To predict the accessibility of KRAS binding sites, we constructed composite systems by aligning crystallographic KRAS with KRAS from all-atom MD simulations (ninety 5 μs simulations of GTP-bound KRAS, excluding the first 2 μs of each simulation). RAF1 proteins in these composite systems are then evaluated for steric overlap with lipids in the membrane associated with the simulated KRAS protein. The number of protein residues that clash with lipids (any protein atom within 0.1 nm of any lipid atom), 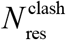, is used to assess the competence of the specific KRAS orientation to bind another protein in the absence of any lipidic accommodation. Orientation-specific ensemble averaging over a total of 270,000 simulation snapshots provides Boltzmann-weighted averages over the sampled displacement of the KRAS G-domain from the center of mass of the lipid bilayer along its normal, *d*_Gz_.

**Supplementary Figure 1.**
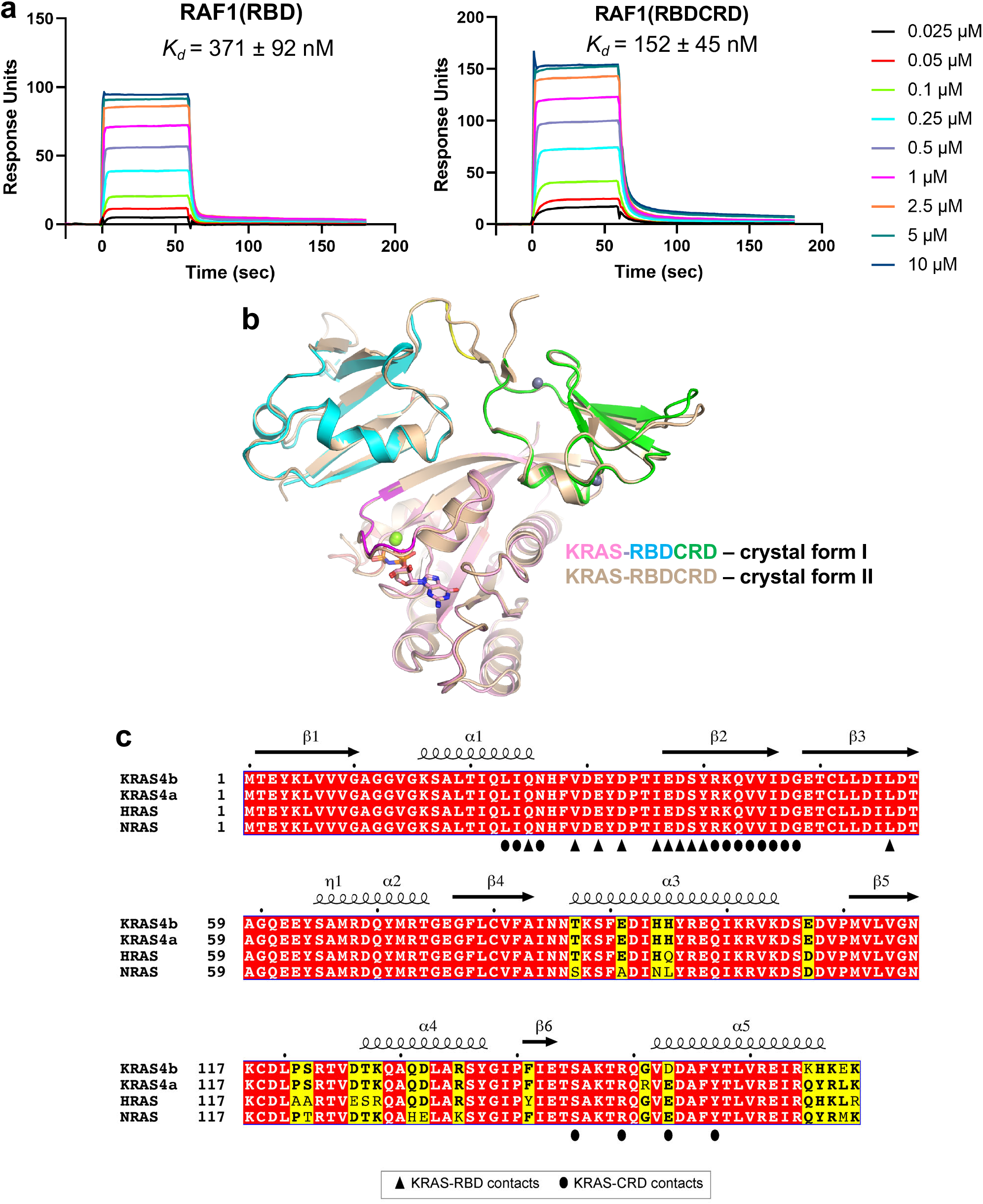
SPR binding kinetics, comparison of two crystal forms of KRAS-RBDCRD complex, and conservation of RBDCRD interacting residues in RAS isoforms. **(a)** SPR sensorgrams showing the binding of RAF1(RBD) and RAF1(RBDCRD) to KRAS. RBD and RBDCRD (0.025 – 10 μM) were injected over avi-tagged WT KRAS, and binding responses were determined. **(b)** Structural superposition of the KRAS-RBDCRD complex solved in two different crystal forms shows a similar arrangement of KRAS and RBDCRD inside the crystal. Crystal form I is colored using the same color scheme as Figure 1c, whereas crystal form II is colored in light brown color. **(c)** Amino acid sequence alignment of the G-domain of human KRAS4b, KRAS4a, HRAS, and NRAS. Fully and partially conserved residues among the RAS isoforms are highlighted in red and yellow, respectively. The secondary structural elements of KRAS seen in the KRAS-RBDCRD structure are shown above the alignment. The KRAS residues that are involved in the interaction with RBD and CRD are indicated below the alignment with upright triangles, and ovals, respectively.

**Supplementary Figure 2.**
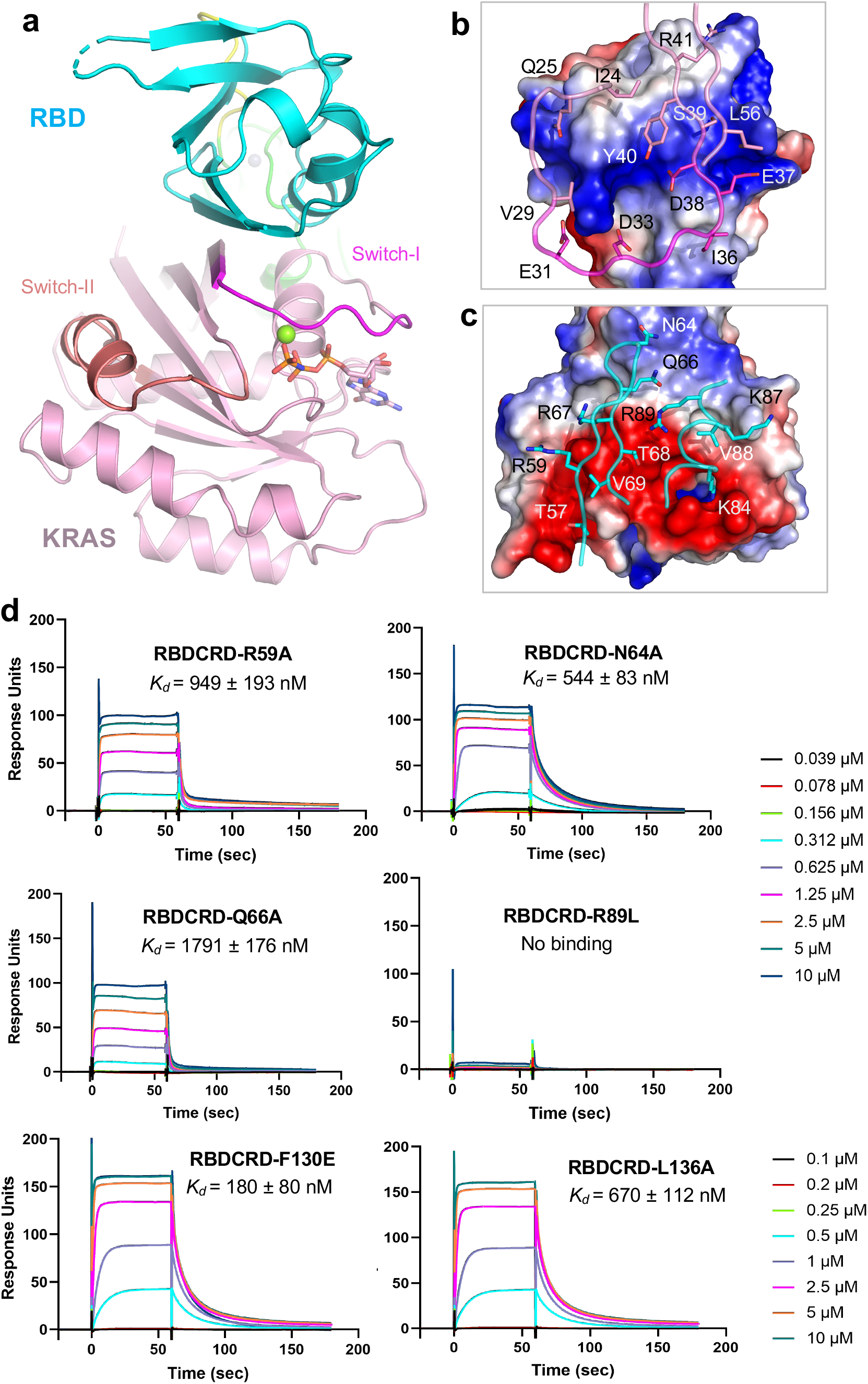
Structural and mutational analysis of KRAS and RBD residues present at the interface in the KRAS-RBDCRD complex. **(a)** The overall structure of the KRAS-RBDCRD complex shown in cartoon representation, highlighting the KRAS-RBD interface. **(b)** The KRAS-RBD interaction interface in the KRAS-RBDCRD structure. RBD is shown in electrostatic surface representation and the KRAS residues that participate at the interface are shown in stick representation. **(c)** The KRAS-RBD interaction interface in the KRAS-RBDCRD structure. KRAS is shown in electrostatic surface representation and the RBD residues that participate at the interface are shown in stick representation. **(d)** SPR sensorgrams showing the binding of RAF1(RBDCRD) mutants to WT KRAS. RBDCRD mutants R59A, N64A, Q66A, and R89L, were injected over avi-tagged WT KRAS in the concentration range (0.039 – 10 μM), and binding responses were determined. RBDCRD mutant F130E and L136A were injected over avi-tagged WT KRAS in the concentration range 0.1 - 20 μM.

**Supplementary Figure 3.**
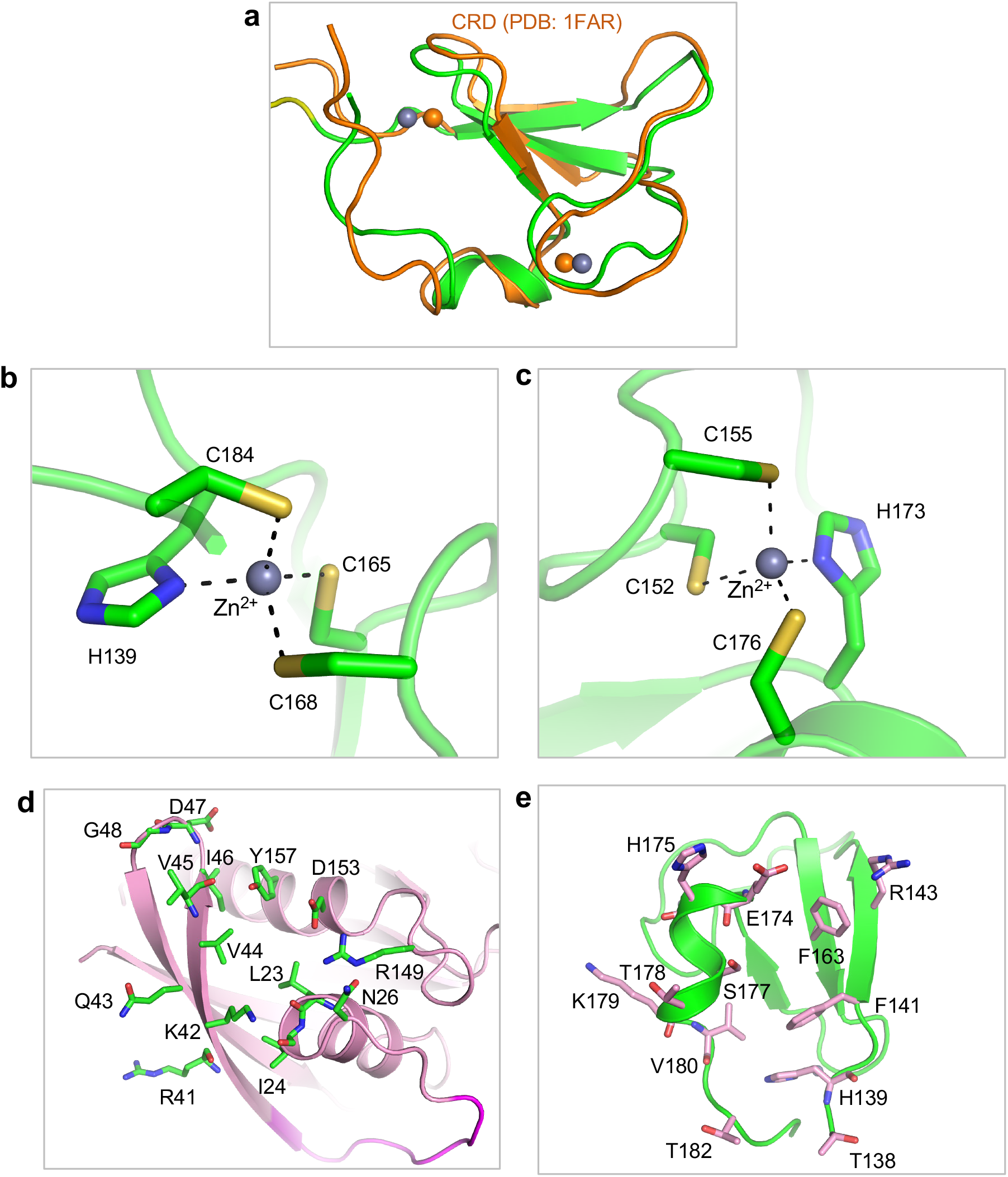
Comparison of NMR and crystal structures of CRD, structural analysis of zinc fingers, and residues present at the KRAS-CRD interface. **(a)** Structural superposition of CRD observed in the KRAS-RBDCRD complex with the NMR (minimized average) structure of CRD (PDB ID: 1FAR) shows overall structural similarity. (**b, c**) Two zinc-finger motifs present in the CRD region of RBDCRD. Zinc(II) ions are shown as spheres, whereas CRD residues are shown in stick representation. Metal coordination bonds are indicated by dashed black lines. **(d)** KRAS (colored pink) is shown in cartoon representation with residues interacting with CRD highlighted in green. Main chains are shown for those residues that partly or solely interact with CRD via their main chain atoms. **(e)** CRD (colored green) is shown in cartoon representation with residues interacting with KRAS highlighted in pink. Main chains are shown for those residues that partly or solely interact with KRAS via their main chain atoms.

**Supplementary Figure 4.**
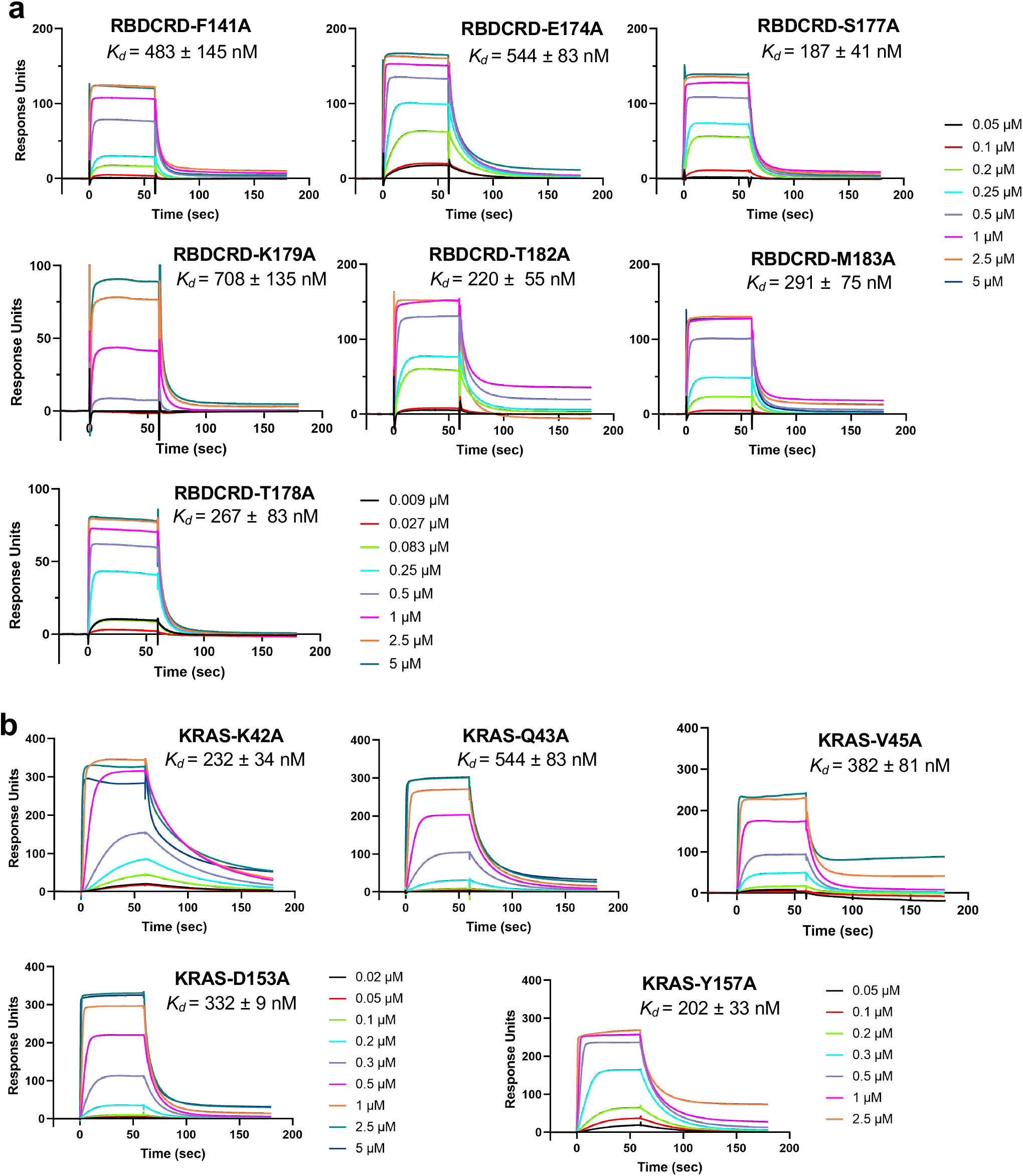
SPR binding analysis of RBDCRD and KRAS mutants present at the KRAS-CRD interface. **(a)** SPR sensorgrams showing the binding of RAF1(RBDCRD) mutants to WT KRAS. RBDCRD mutants F141A, E174A, S177A, T178A, K179A, T182A, and M183A, were injected over avi-tagged WT KRAS in the concentration range (0.009 – 5 μM), and binding responses were determined. **(b)** SPR sensorgrams showing the binding of KRAS mutants to WT RBDCRD. WT RBDCRD was injected over avi-tagged KRAS mutant in the concentration range (0.02 - 5 μM), and binding responses were determined. For the KRAS mutant Y157A, WT RBDCRD was injected over avi-tagged KRAS-Y157A in the concentration range 0.05 - 2.5 μM.

**Supplementary Figure 5.**
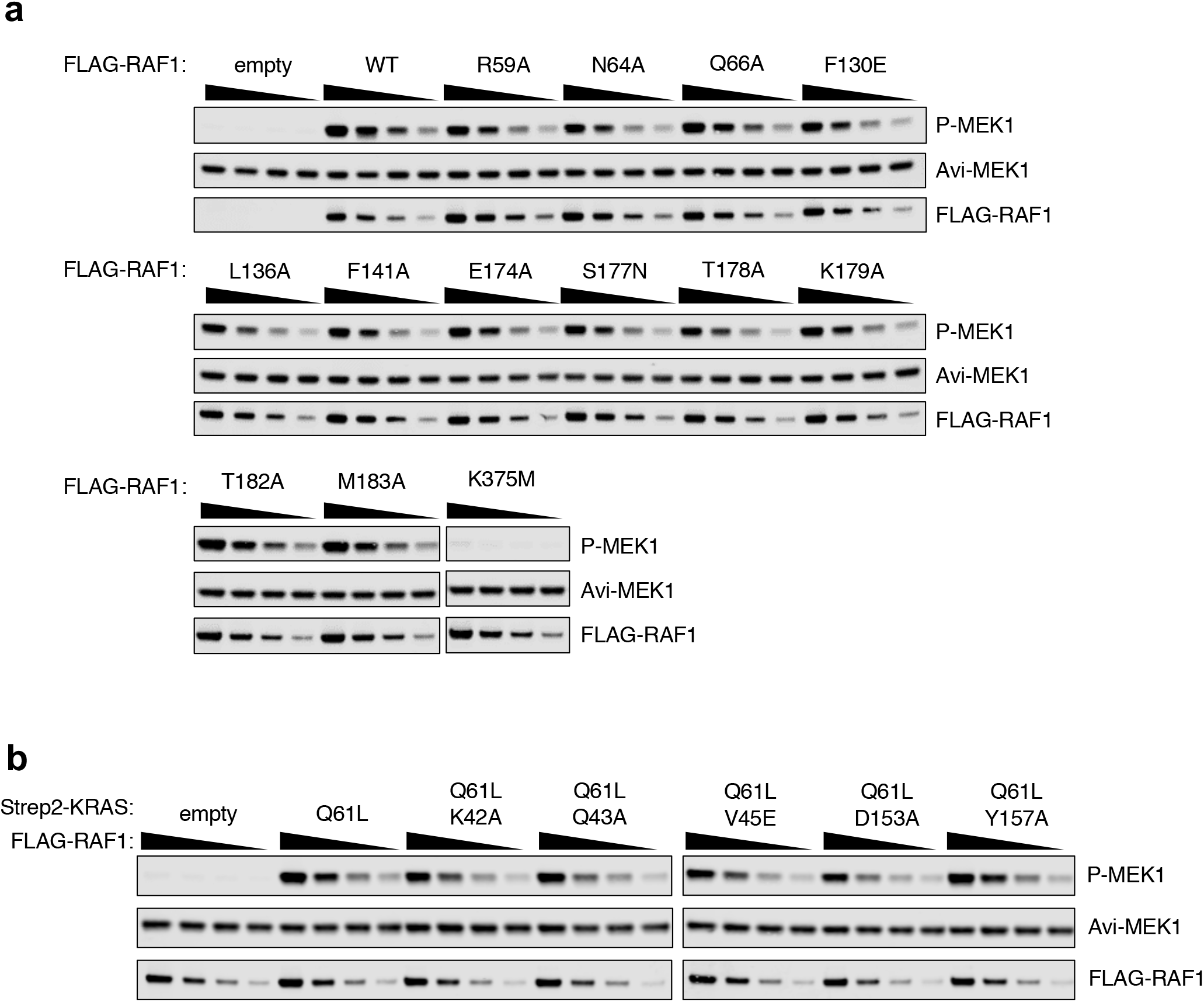
Western blots showing the effects of point mutations at the KRAS-RBDCRD interface on KRAS-RAF1 binding in 293T cells and RAF1 kinase activity. **(a)** Serial dilutions of WT and point mutants of RAF1 (purified from FLAG IPs in Fig. 5a) were used to monitor kinase activity towards recombinant MEK1. **(b)** Kinase activity of RAF1 purified from FLAG IPs in Fig. 5c was measured as described in panel a.

**Supplementary Figure 6.**
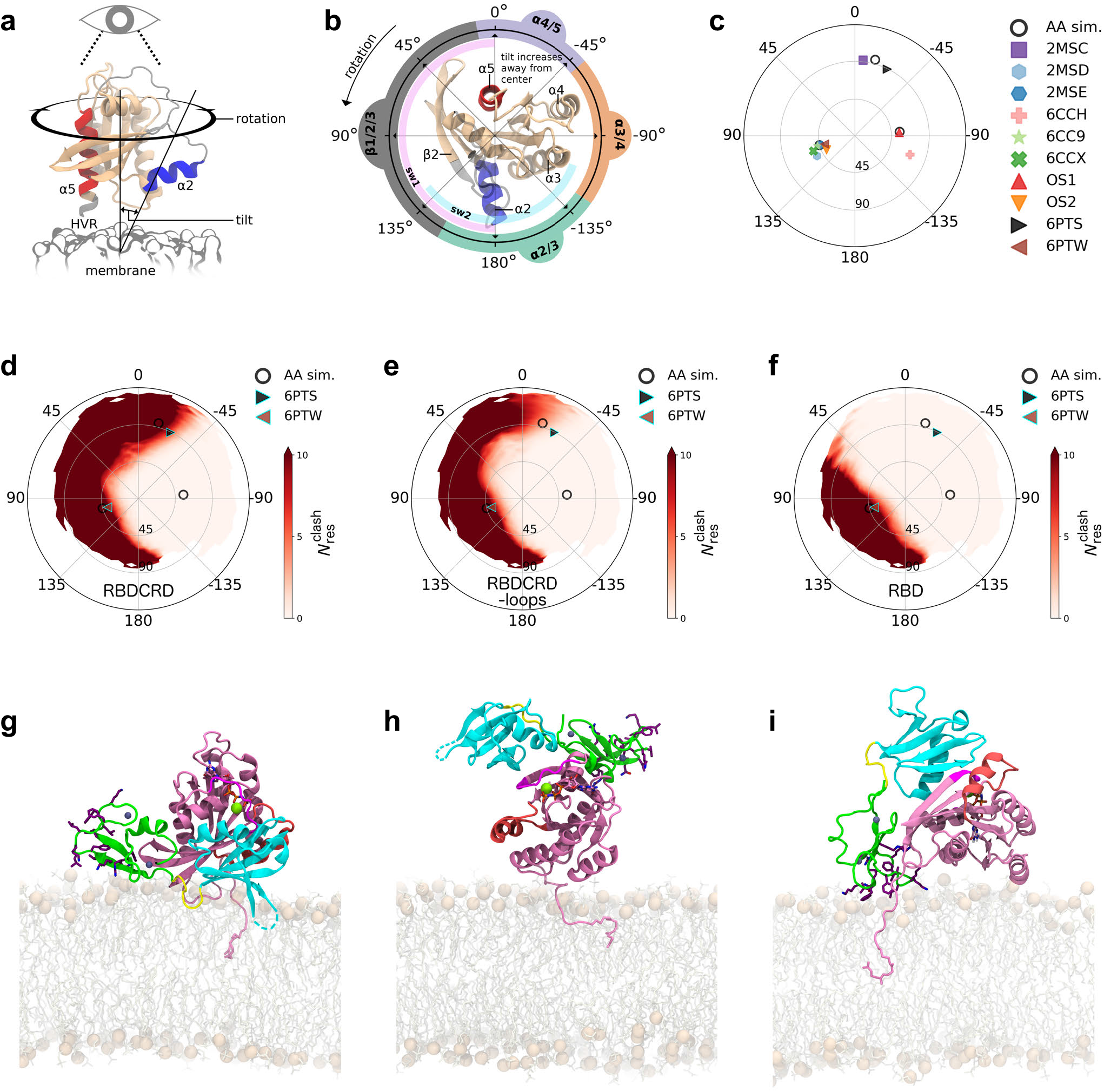
Orientation of KRAS in relation to the membrane surface and occlusion of RAF1 binding. (**a, b**) Definition of tilt and rotation angles. Regions of KRAS that approach the membrane upon tilting are inscribed around the perimeter of panel b. Reprinted with permission from *Biophysical Journal*^28^. (**c**) Reported membrane orientations of KRAS. “AA sim.” are from simulations of KRAS on mixed anionic: zwitterionic lipid bilayers^28^. PDB: 2MSC, 2MSD, and 2MSE are the “exposed,” “occluded,” and “semi-occluded” orientations from Mazhab-Jafari *et al*.^52^; PDB: 6CCH, 6CC9, and 6CCX are the “E3,” “O1,” and “O2” orientations from Fang *et al*.^53^ “OS1” and “OS2” are from simulations by Prakash *et al*.^54^. PDB: 6PTS and 6PTW are the “state A” and “state B” orientations of KRAS-RBDCRD from Fang *et al*.^6^. Ensemble average orientations are reported for PDB entries comprising multiple models. (**d-f**) Relationship between KRAS orientation and steric competence to bind (d) RBD, (e) RBDCRD, and (f) RBDCRD excluding the CRD’s mixed hydrophobic/cationic loop residues R_143_KTFLKLAF_151_ and K_157_FLLNGFR_164_. 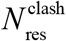 is the number of RAF1 residues that overlap with membrane lipids (any protein atom within 0.1 nm of any lipid atom). Relatively large values of, shown as red in panels d-f, indicate occlusion of protein-protein interactions by approximately planar models of the cell membrane. In some cases, this occlusion may be relieved by structural accommodation of membrane lipids. The difference between panels e and f represents evaluation of occlusion according to models in which hydrophobic CRD loops are (e) disallowed or (f) allowed to penetrate the membrane surface. (**g-h**) Additional models of membrane interactions of the crystallographic RAS-RBDCRD complex, analogous to Fig. 7c except orienting KRAS such that (g) β sheets β1-3 and switch-I or (h) a helices a3 and a4 face the membrane. (**i)** Similar to Fig. 7c, except KRAS and RAF1 are oriented and configured as in model 0 of PDB: 6PTS^29^.

**Supplementary Table 1:**
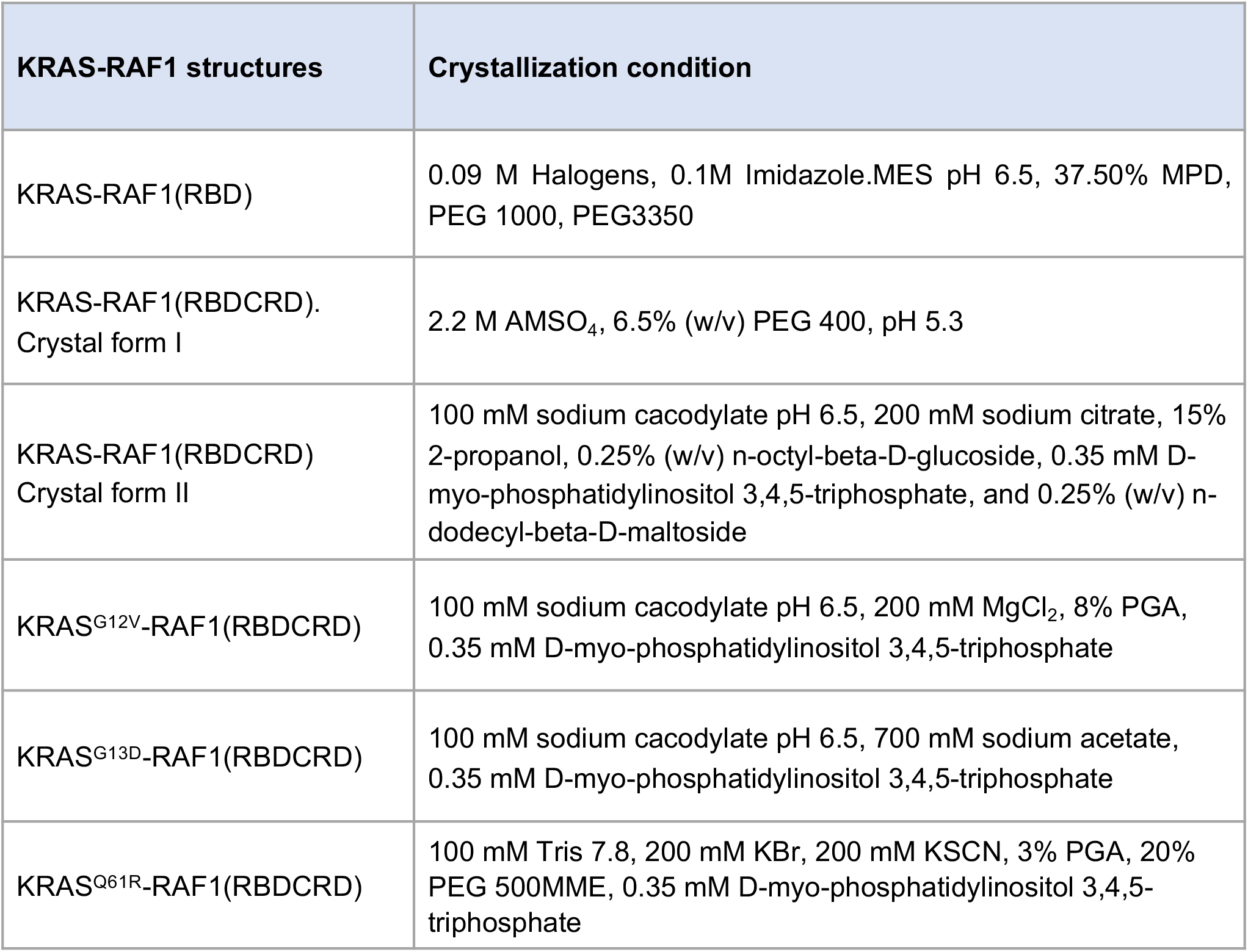
Crystallization conditions for the structures described in this study.

**Supplementary Table 2:**
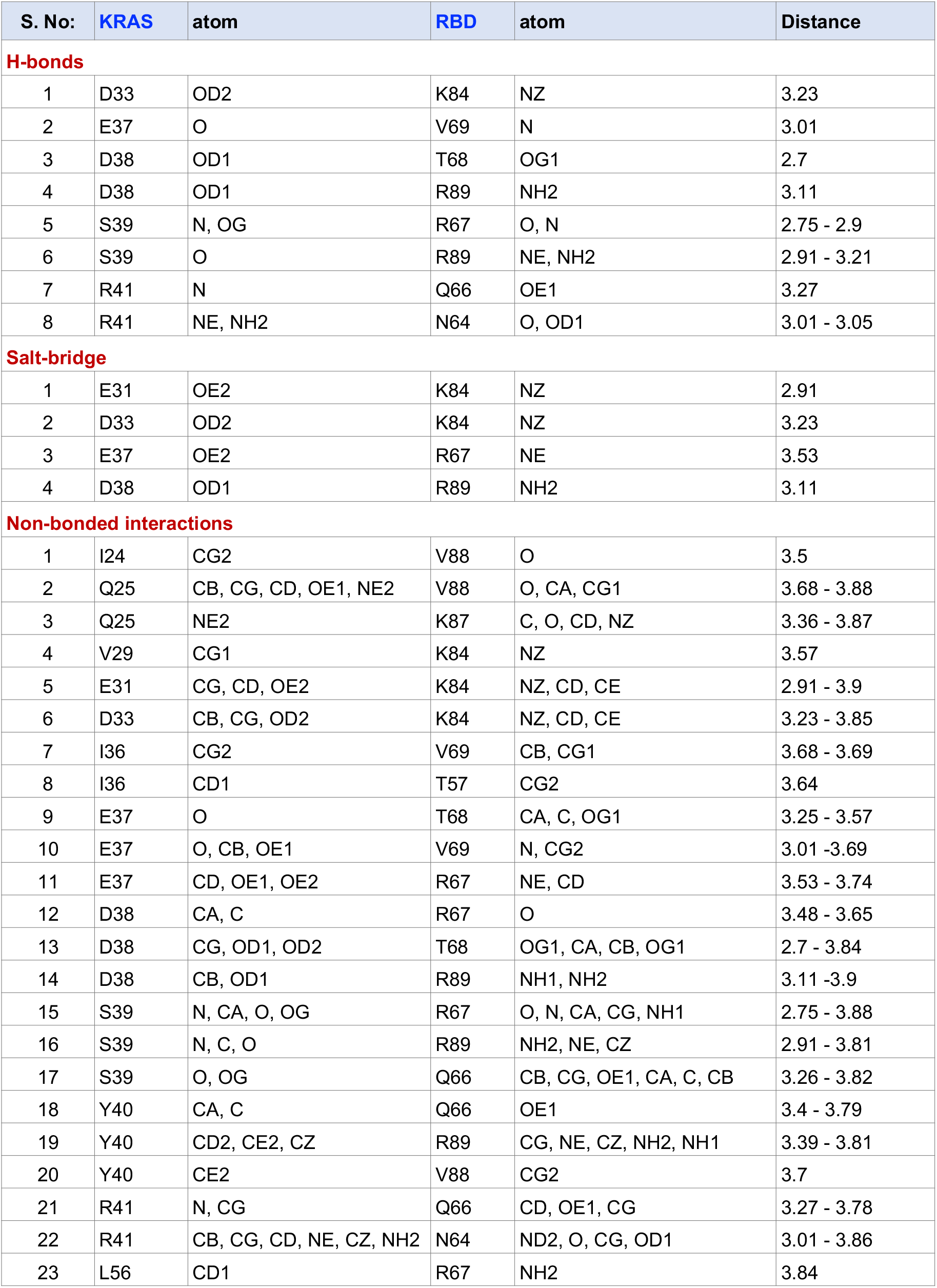
List of protein-protein interactions present at the KRAS-RAF1(RBD) interface in the structure of KRAS-RAF1(RBDCRD) complex.

**Supplementary Table 3:** List of protein-protein interactions present at the KRAS-RAF1(RBD) interface in the structure of KRAS-RAF1(RBD) complex.

| S. No: | KRAS | atom | RBD | atom | Distance |
| --- | --- | --- | --- | --- | --- |
| **H-bonds** |  |  |  |  |  |
| 1 | E31 | OD2 | K84 | NZ | 2.77 |
| 2 | E37 | O | V69 | N | 3.11 |
| 3 | E37 | OE1, OE2 | R67 | NE, NH2 | 2.74 - 3.07 |
| 4 | E37 | OE2 | R59 | NE, NH2 | 2.64 - 2.82 |
| 5 | D38 | OD1 | T68 | OG1 | 2.61 |
| 6 | D38 | OD1 | R89 | NH2 | 2.91 |
| 7 | S39 | N, OG | R67 | O, N | 2.91 - 2.94 |
| 8 | S39 | O | R89 | NE, NH2 | 2.89 - 3.07 |
| 9 | R41 | N | Q66 | OE1 | 3.11 |
| 10 | R41 | NH1 | N64 | OD1 | 2.95 |
| **Salt-bridge** |  |  |  |  |  |
| 1 | E31 | OE2 | K84 | NZ | 2.77 |
| 2 | E37 | OE2 | R59 | NE | 2.64 |
| 3 | E37 | OE1 | R67 | NE | 2.74 |
| 4 | D38 | OD1 | R89 | NH2 | 2.91 |
| **Non-bonded interactions** |  |  |  |  |  |
| 1 | I24 | CG2 | V88 | O | 3.31 |
| 2 | Q25 | OE1 | V88 | CG1 | 3.71 |
| 3 | E31 | CG, CD, OE2 | K84 | NZ, CE | 2.77 - 3.55 |
| 4 | I36 | CB, CD1 | V69 | CG1 | 3.89 - 3.9 |
| 5 | I36 | CD1 | T57 | CG2 | 3.89 |
| 6 | E37 | O | T68 | CA, C | 3.42 - 3.63 |
| 7 | E37 | O, CG, OE2 | V69 | N, CG2 | 3.11 - 3.81 |
| 8 | E37 | CD, OE2 | R59 | NE, NH2, CD, CZ | 2.64 - 3.84 |
| 9 | E37 | CD, OE1, OE2 | R67 | NE, NH2, CB, CD, CZ | 2.74 - 3.88 |
| 10 | D38 | CA, C, OD1 | R67 | O | 3.33 - 3.84 |
| 11 | D38 | CG, OD1, OD2 | T68 | OG1, CA, CB, OG1 | 2.61 - 3.67 |
| 12 | D38 | CG, OD1 | R89 | NH1, CZ, NH2 | 2.91 - 3.86 |
| 13 | S39 | N, O, OG | R67 | O, N, CA, CB | 2.91 - 3.83 |
| 14 | S39 | N, C, O | R89 | NH2, NE, CZ | 2.89 - 3.78 |
| 15 | S39 | O, OG | Q66 | CB, CG, CA, C | 3.41 - 3.6 |
| 16 | Y40 | CA, C | Q66 | OE1 | 3.31 - 3.64 |
| 17 | Y40 | CD2, CE2, CZ, OH | R89 | CG, NE, CZ, NH2, NH1 | 3.37 - 3.84 |
| 18 | Y40 | CE2 | V88 | CG2 | 3.72 |
| 19 | R41 | N | Q66 | CD, OE1 | 3.11 - 3.8 |
| 20 | R41 | CB, CG, CD, NH1 | N64 | ND2, O, CG, OD1 | 2.95 - 3.84 |
| 21 | R41 | CZ, NH1, NH2 | K65 | CD | 3.53 - 3.77 |
| 22 | M67 | SD | R59 | NH2 | 3.89 |

**Supplementary Table 4:**
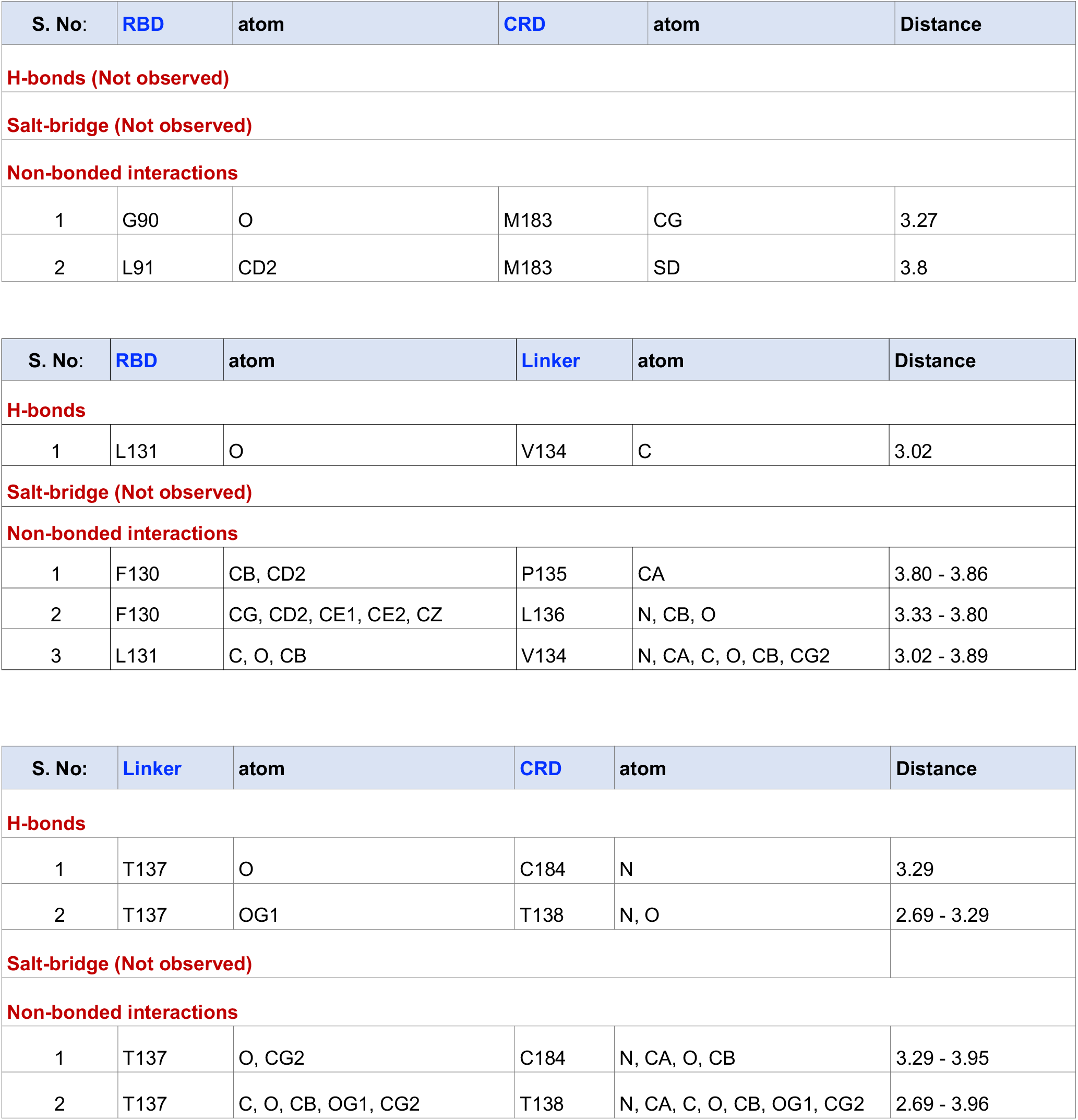
List of interdomain interactions present at the RBD-Linker-CRD interface in the structure of KRAS-RAF1(RBDCRD) complex.

| S. No: | RBD | atom | CRD | atom | Distance |
| --- | --- | --- | --- | --- | --- |
| **H-bonds (Not observed)** |  |  |  |  |  |
| **Salt-bridge (Not observed)** |  |  |  |  |  |
| **Non-bonded interactions** |  |  |  |  |  |
| 1 | G90 | O | M183 | CG | 3.27 |
| 2 | L91 | CD2 | M183 | SD | 3.8 |

| S. No: | RBD | atom | Linker | atom | Distance |
| --- | --- | --- | --- | --- | --- |
| **H-bonds** |  |  |  |  |  |
| 1 | L131 | O | V134 | C | 3.02 |
| **Salt-bridge (Not observed)** |  |  |  |  |  |
| **Non-bonded interactions** |  |  |  |  |  |
| 1 | F130 | CB, CD2 | P135 | CA | 3.80 - 3.86 |
| 2 | F130 | CG, CD2, CE1, CE2, CZ | L136 | N, CB, O | 3.33 - 3.80 |
| 3 | L131 | C, O, CB | V134 | N, CA, C, O, CB, CG2 | 3.02 - 3.89 |

| S. No: | Linker | atom | CRD | atom | Distance |
| --- | --- | --- | --- | --- | --- |
| **H-bonds** |  |  |  |  |  |
| 1 | T137 | O | C184 | N | 3.29 |
| 2 | T137 | OG1 | T138 | N, O | 2.69 - 3.29 |
| **Salt-bridge (Not observed)** |  |  |  |  |  |
| **Non-bonded interactions** |  |  |  |  |  |
| 1 | T137 | O, CG2 | C184 | N, CA, O, CB | 3.29 - 3.95 |
| 2 | T137 | C, O, CB, OG1, CG2 | T138 | N, CA, C, O, CB, OG1, CG2 | 2.69 - 3.96 |

**Supplementary Table 5:** List of protein-protein interactions present at the KRAS-RAF1(CRD) interface in the structure of KRAS-RAF1(RBDCRD) complex.

| S. No: | KRAS | atom | CRD | atom | Distance |
| --- | --- | --- | --- | --- | --- |
| **H-bonds** |  |  |  |  |  |
| 1 | K42 | NZ | V180 | O | 2.67 |
| 2 | Q43 | NE2 | H139 | O | 3.03 |
| 3 | V45 | N | S177 | OG | 3.09 |
| 4 | D47 | N | E174 | OE2 | 2.74 |
| 5 | G48 | N | E174 | OE1 | 2.59 |
| 6 | G48 | O | R143 | NH1 | 2.76 |
| 7 | R149 | NH2 | T178 | OG1 | 3.1 |
| 8 | D153 | OD1 | T178 | OG1 | 2.9 |
| **Salt-bridge (Not observed)** |  |  |  |  |  |
| **Non-bonded interactions** |  |  |  |  |  |
| 1 | L23 | O, CD1 | T178 | O, CG2 | 3.19 - 3.81 |
| 2 | I24 | O | T182 | OG1 | 3.57 |
| 3 | N26 | CG, OD1, ND2 | K179 | CG, CD, CE | 3.61 - 3.87 |
| 4 | R41 | O | T182 | CG2 | 3.73 |
| 5 | K42 | CD, NZ | T182 | OG1 | 3.53 - 3.82 |
| 6 | K42 | CE, NZ | T178 | O | 3.41 - 3.48 |
| 7 | K42 | CE | V180 | O, C | 3.57 - 3.86 |
| 8 | K42 | NZ | V180 | O | 2.67 |
| 9 | Q43 | CD, OE1, NE2 | H139 | O | 3.03 - 3.63 |
| 10 | Q43 | OE1 | F141 | CD2 | 3.72 |
| 11 | Q43 | NE2 | T138 | CB | 3.84 |
| 12 | V44 | CG1 | S177 | OG | 3.58 |
| 13 | V44 | CG1 | T178 | CG2 | 3.53 |
| 14 | V45 | N, CA, O, CB | S177 | CB, OG | 3.09 - 3.85 |
| 15 | V45 | C, O | E174 | CG, O, CD | 3.18 - 3.83 |
| 16 | V45 | CG1 | F163 | CE1, CZ | 3.45 - 3.53 |
| 17 | V45 | CG1 | E174 | CG, OE1 | 3.57 - 3.87 |
| 18 | I46 | C | E174 | OE2 | 3.75 |
| 19 | D47 | N, CA, C | E174 | CD, OE1, OE2 | 2.74 - 3.85 |
| 20 | G48 | N, CA | E174 | CD, OE1, OE2 | 2.59 - 3.67 |
| 21 | G48 | CA, C, O | R143 | NH1, CZ | 3.65 - 3.87 |
| 22 | G48 | O | F163 | CZ | 3.78 |
| 23 | R149 | CZ, NH1 | T178 | OG1, CB | 3.1 - 3.84 |
| 24 | R149 | NH1 | K179 | CE | 3.78 |
| 25 | D153 | CG, OD1 | T178 | OG1, CG2, CB | 2.9 - 3.88 |
| 26 | D153 | OD2 | H175 | O | 3.89 |
| 27 | D153 | OD2 | T178 | OG1 | 3.09 |
| 28 | Y157 | CE1 | T178 | CG2 | 3.76 |
| 29 | Y157 | OH | E174 | O | 3.4 |
| 30 | Y157 | OH | S177 | OG | 3.48 |

